# Reliable ligand discrimination in stochastic multistep kinetic proofreading: First passage time vs. product counting strategies

**DOI:** 10.1101/2024.02.06.579249

**Authors:** Xiangting Li, Tom Chou

## Abstract

Cellular signaling, crucial for biological processes like immune response and homeostasis, relies on specificity and fidelity in signal transduction to accurately respond to stimuli amidst biological noise. Kinetic proofreading (KPR) is a key mechanism enhancing signaling specificity through time-delayed steps, although its effectiveness is debated due to intrinsic noise potentially reducing signal fidelity. In this study, we reformulate the theory of kinetic proofreading (KPR) by convolving multiple intermediate states into a single state and then define an overall “processing” time required to traverse these states. This simplification allows us to succinctly describe kinetic proofreading in terms of a single waiting time parameter, facilitating a more direct evaluation and comparison of KPR performance across different biological contexts such as DNA replication and T cell receptor (TCR) signaling. We find that loss of fidelity for longer proofreading steps relies on the specific strategy of information extraction and show that in the first-passage time (FPT) discrimination strategy, longer proofreading steps can exponentially improve the accuracy of KPR at the cost of speed. Thus, KPR can still be an effective discrimination mechanism in the high noise regime. However, in a product concentration-based discrimination strategy, longer proofreading steps do not necessarily lead to an increase in performance. However, by introducing activation thresholds on product concentrations, can we decompose the product-based strategy into a series of FPT-based strategies to better resolve the subtleties of KPR-mediated product discrimination. Our findings underscore the importance of understanding KPR in the context of how information is extracted and processed in the cell.

**Author summary:** Kinetic proofreading (KPR) is mechanism often employed by cells to enhance specificity of ligand-receptor. However, the performance of kinetic proofreading may be hampered by noise and a low signal-to-noise ratio. By consolidating multiple kinetic proofreading steps into a single state and assigning an associated waiting, or “processing time,” we developed an analytic approach to quantify the performance of KPR in different biological contexts. Despite a trade-off between speed and accuracy inherent to a first-passage time KPR strategy, we show that a signaling molecule-based discrimination strategy can enhance the performance benefits of KPR. We further decompose the product-based discrimination strategy into a set of first-passage times to different thresholds of signaling molecules produced. Through this decomposition, we find that a threshold that adjusts dynamically throughout the recognition process depends on the duration of the process. We propose that this more nuanced product-based KPR-mediated recognition process can be realized biologically. The precise structural basis for a dynamic threshold merits further experimental exploration, as it may hold significant implications for understanding biological mechanisms of information transmission at a molecular level.

## Introduction

Various cellular processes require a high degree of specificity in order to function properly, including DNA replication, gene expression, and cellular signaling. The degree of specificity observed is often hard to justify by a simple binding-affinity argument, the specificity of which is proportional to exp(*−*ΔΔ*G/RT*), where ΔΔ*G* is the difference in free energy between the correct and incorrect ligands [1]. For example, the estimated error probability per nucleotide in DNA replication is estimated to be 10^−9^ [2], but the net free energy difference between mismatched and matched base pairs is only 0.5 kcal*/*mol [3], suggesting the theoretical error rate would be *∼* 10^−3^. Similarly, T cells need to specifically distinguish self-antigens from mutant self-antigens, also known as neoantigens, which can differ by only one or a few amino-acids [4].

Kinetic proofreading (KPR) [1, 5, 6] typically denotes a chemical reaction mechanism that can significantly increase the specificity towards a desired ligand against competing ligands. In the KPR context, “proofreading” is accomplished by introducing additional irreversible, energy-consuming kinetic steps which individually may not distinguish desired ligands from undesired ones. However, these steps impart a delay to final product release allowing for “resetting” of the process and an overall lower final error rate (a multistep KPR mechanism is illustrated in Fig. 1(a) below and mathematical details are discussed in the Materials and Methods).

**Fig 1.**
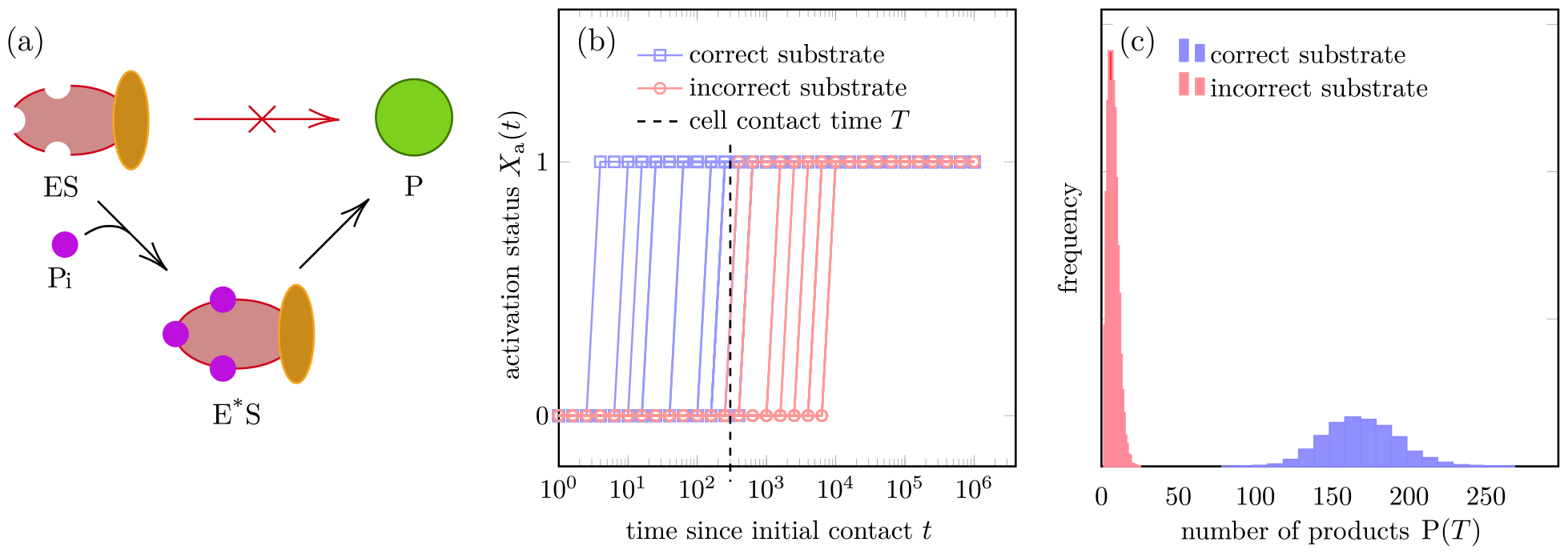
Schematic of the KPR process and different strategies of interpreting the output. (a) The complex of enzyme and substrate ES alone cannot produce the final product P. It has to undergo a number of proofreading steps, represented here by phosphorylation (Pi), before the activated state E*S can produce the final product. (b) The FPT-based discrimination strategy, simply reaching the activated state E*S is interpreted as the output. *X*_a_(*t*) = 0 if the system has not reached the activated state by time *t*, and *X*_a_(*t*) = 1 otherwise. At *t* = 0, the reaction starts with the enzyme in the free state E. The dashed vertical line represents the termination of entire process, *e.g*., the time *T* at which the T cell and antigen presenting cell (APC) separate. *X*_*a*_(*t*) = 1 for some *t < T* is interpreted as a positive response. (c) The product-based discrimination strategy. In this strategy, the number of product molecules P(*T*) produced within a given time *T* is interpreted as the output. Due to the intrinsic noise, the number of product molecules is a random variable and their distribution is shown for correct and incorrect ligands.

First proposed by Hopfield to explain the high specificity of DNA/protein synthesis [1], the KPR mechanisms have been invoked to explain other biological processes such as T cell receptor (TCR) signaling [6] and microtubule growth [7]. These first treatments of KPR described it within the steadystate limit of deterministic mass-action models, comparing the steady-state fluxes of the correct and incorrect product formation. Reactions inside the cell, however, are often between small numbers of molecules and are thus stochastic. Stochastic aspects of KPR have also been considered, emphasizing the statistics of first passage times (FPT) to product formation [8–10].

Recently, Kirby and Zilman reported that adding more kinetic proofreading steps almost always decreases the signal-to-noise ratio (SNR) defined by the ratio of the mean to the standard deviation of the output signal (the number of signaling molecules produced), suggesting that KPR is not an optimal strategy for information processing due to noise [11]. However, TCR signaling and T cell activation, the context that Kirby and Zilman describe, is a highly specific process that does involve multiple kinetic proofreading steps, but with adaptive variants of KPR used to model TCR signaling [12–16].

In this paper, we reconcile the apparent contradiction between the high specificity of TCR signaling and the low SNR of a longer-chain KPR process. The key theoretical insight involves convolving the multiple intermediate irreversible steps into a single equivalent state in which the system stays for time *τ*. Instead of explicitly treating a series of sequential states, we define a single, equivalent waiting time or “processing” time *τ*. This simple reduction reveals intriguing insights, allows us to analytically and systematically explore different biological contexts of KPR, and provides an easier framework on which to test different metrics of KPR performance.

We show that the apparent contradiction arises from different strategies of determining whether a final output is correct. In the FPT-based scenario for both DNA replication and TCR signaling, arriving at an activated state or a “product” state within a given time is interpreted as the output, as illustrated in Fig. 1(b). A longer processing time *τ* can exponentially improve the accuracy of KPR at the cost of speed. The trade-off between speed and accuracy has been reported in experimental studies [17–19]. In an alternative strategy for TCR signaling implicitly used by Kirby and Zilman, the maximum SNR, or mutual information between the input and output arises with just two proofreading steps. However, more processing steps leads to a decrease in KPR performance.

The alternative discrimination strategy implicit in Kirby’s metric is to detect the number of products (*e.g*., signaling molecules that lead to downstream processes) generated within a finite time without explicitly resolving the final response of the cell, as is illustrated in Fig. 1(c). Mutual information has been used in recent studies to quantify the information flow in cellular decision-making processes [20–22]. Here, we introduce mutual information and channel capacity in order to compare the performance of the two strategies (FPT to a target state and product counting) on equal footing. We also construct a decomposition of the product-based strategy into a series of FPT-based strategies with different product molecule thresholds and conclude that the product detection strategy is equivalent to a strategy that dynamically adjusts the threshold according to the duration of the process. This dynamic thresholding strategy can be shown to be more robust to fluctuations over the duration of the reaction. The effectiveness of this strategy can be attributed to an additional layer of proofreading.

Our analysis and findings present a unified framework for analyzing KPR under different biological scenarios. We also highlight the importance of understanding how different strategies of information extraction can affect the performance and parameter tuning of KPR.

## Materials and methods

In this section, we first describe the general model of KPR and then apply it to two specific biological contexts, DNA replication and TCR signaling.

### Model settings

In the conventional setting of KPR, the complex E^(0)^S composed of receptor E and a “correct” substrate (or ligand) S forms and dissociates with binding and unbinding rates *k*_*±*1_. A complex with the “incorrect” ligand forms and dissociates with rates *q*_*±*1_. Both complexes can undergo multiple nonequilibrium transitions or proofreading steps (*e.g*., sequential phosphorylation) traversing internal states (E^(1)^S, …, E^(*m−*1)^S) before the final product P can be released or produced by the fully activated state E*S.

Each internal state of the complex can dissociate with rate *k*_off_ or proceed to the next step with rate *k*_f_, as shown in Fig. 2(a). To simplify our subsequent analysis, we set *k*_off_ = *k*_*−*1_ as in [11]. Additionally, in the multistep limit (*m → ∞*), the processing time *τ* of reaching the final activated state E*\** S from ES is fixed to a deterministic value *τ* = *m/k*_f_, (in which *k*_f_ scales with *m* as *m → ∞*) as was analyzed by [9]. We can thus simplify the reaction diagram in Fig. 2(a) by lumping the internal states (E^(0)^S, …, E^(*m−*1)^S) into a single state ES as shown in Fig. 2(b). Note that because of independence between activation and dissociation in these steps, the transition of the aggregated state ES to the dissociated state E + S is still Markovian with the same rate *k*_*−*1_. The master equation of the simplified model and its relation to the original multi-step model are discussed in Appendix A1.

**Fig 2.**
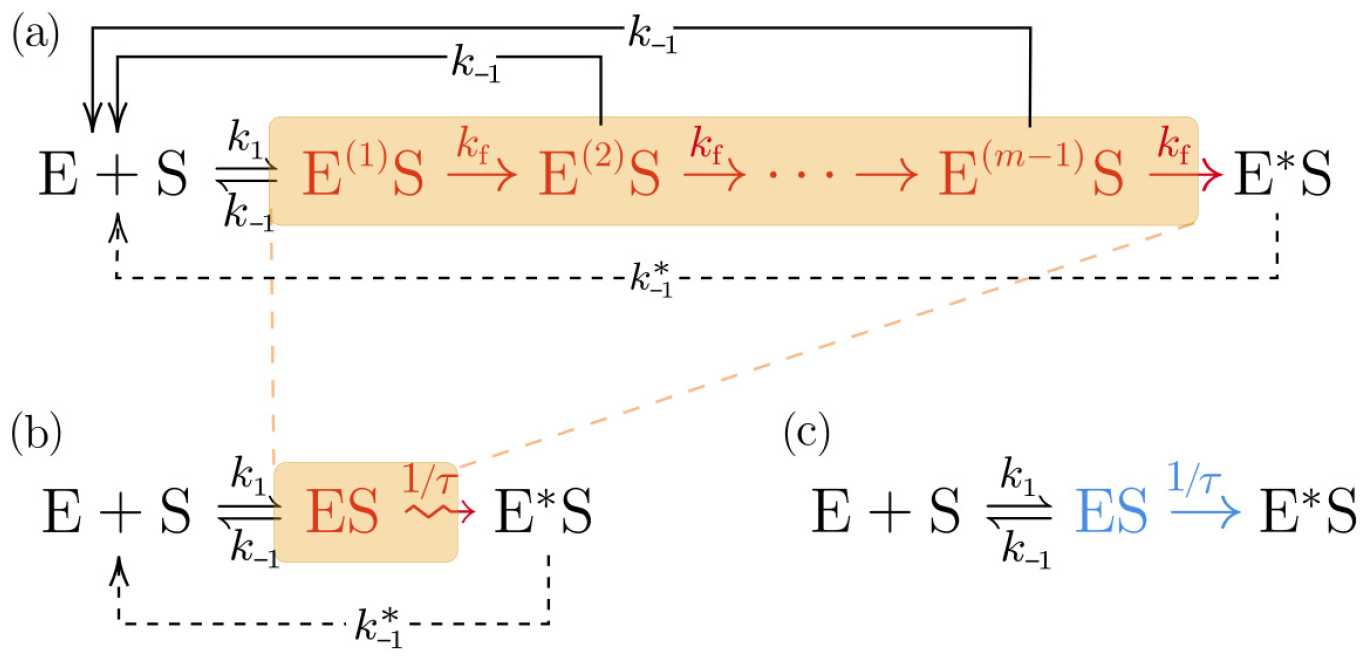
Reaction schemes of different descriptions of the simple kinetic proofreading process. (a) The conventional description of KPR explicitly incorporating multiple proofreading steps. (b) A reduced representation of KPR in which multiple driven steps are lumped in a single proofreading step. For comparison, we show the classical Michaelis-Menten reaction scheme in (c). In (a) and (b), unbinding of E*S is not essential, *i.e*., it is possible that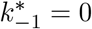, thus marked in dashed lines. Additionally, in (a-b), there can be a production step after E*S is reached, which is not explicitly shown.

As shown in Fig. 2(c), our simplified model has a similar structure to the classical Michaelis-Menten reaction scheme. However, our simplified model captures a crucial element of the KPR mechanism. In the Michaelis-Menten scheme, the waiting time *τ* in the complex state ES before converting to product is exponentially distributed. In the KPR scheme, the waiting time *τ* in the state ES before converting to the activated state E*S is assumed to be non-exponentially distributed. An exponentially distributed waiting time reflects a memoryless process in which the evolution of the system depends only on the current state. This memoryless property is the defining feature of a Markov process. By contrast, memory in the KPR reaction process results in a non-exponential distribution of the processing time *τ*. In the simplification of the KPR we are considering, the processing time is fixed to the value *τ, i.e*., its distribution is a Dirac delta function at *τ*. For example, the waiting time of activation does not depend only on the current state, but also on the time elapsed since the initial formation of the complex, which happened in the past. This non-Markovian step acts as a memory of the system or a clock that keeps track of the time elapsed since the initial formation of the complex, and can be achieved by, *e.g*., tracking the phosphorylation state of the complex ES. The biological context in which KPR operates will be a determining factor in KPR model structure and in the specification the most appropriate performance metric.

### DNA replication setting

First, we consider the DNA replication scenario, specifically a single nucleotide incorporation step catalyzed by DNA polymerase (*E*). We track the system starting from an initial state where the enzyme is free. The correct substrate S refers to the complementary nucleotide to the template strand, while the incorrect substrate S′ refers to the other three nucleotides. The enzyme can bind to either substrate with rates *k*_1_ and *q*_1_, respectively. The enzyme can also unbind from either substrate with rates *k*_*−*1_ and *q*_*−*1_.

The reaction diagram in Fig. 2(b) is translated as

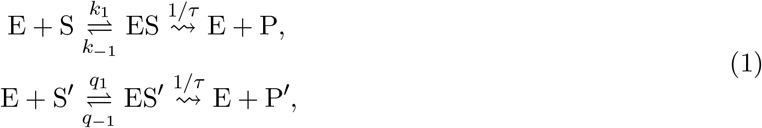

where S and S′ denote the correct and incorrect substrates and P and P′ denote the correct and incorrect products. For simplicity, we identify the activated state E*S of E*S′ with final products E + P or E + P′, referring to incorporation of the correct or incorrect nucleotide, respectively.

We track the system until either one of the products is produced, allowing repeated binding and unbinding of the enzyme to the substrates. While there can be multiple replication forks in a cell, we focus on a single DNA polymerase in this study. Thus, we can represent the stochastic system by a simplified stochastic process with described by three transient states indicating the status of the DNA polymerase; namely, unbound polymerase (E), polymerase bound to correct substrate (ES), and polymerase bound to incorrect substrate (ES′). There are also two absorbing states, namely, correct product (E + P) and incorrect product (E + P′). This simplified stochastic scheme is shown in Fig. 3.

**Fig 3.**
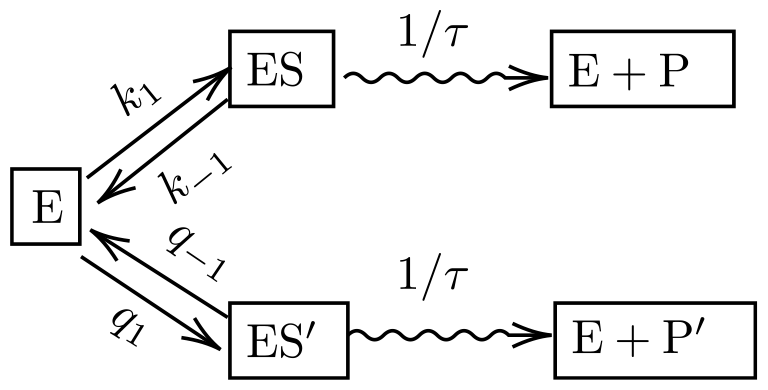
The simplified model of KPR in DNA replication with only one enzyme. Here, the enzyme (*e.g*., DNA polymerase) has three states, namely, free (E), bound to correct substrate (ES), and bound to incorrect substrate (ES′). These states interconvert with rates specified in the model. When the enzyme is bound to substrate, it produces the product (P or P′) after a waiting time *τ*.

The performance of the KPR process is quantified by the probability 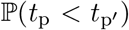 that the FPT *t*_p_ to the correct product state (E + P) is less than the FPT 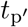 to the wrong product (E + P′). By the standard approach of conditioning on the next state, we find

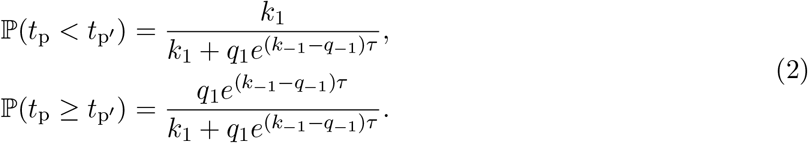

Using the same conditioning approach, we can also calculate the mean first passage time (MFPT) 𝔼[*t*] to either product starting from the free state E:

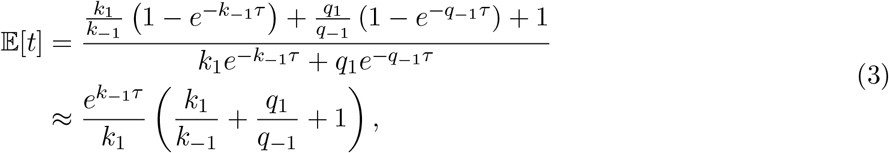

where the above approximation holds if 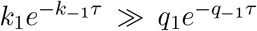, which is typically the case in DNA replication, as 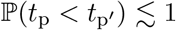.

### TCR signaling setting

Next, we consider proofreading in the TCR signaling context. T cells form contact with an antigen presenting cell (APC) and scans the surface of the APC for the presence of a foreign antigen (or epitope) (S). The TCR on the T cell membrane can bind to the epitope on the APC membrane which initiates a signaling cascade that leads to T cell activation. The activated T cell can then produce signaling molecules that trigger the immune response. If the APC does not carry the foreign antigen corresponding to the TCR, the T cell should not be activated within the contact time *T* and disengages from the APC. There are three features of TCR signaling that are distinct from the DNA signaling process. First, the APC is likely not to carry a foreign antigen, *i.e*., the correct substrate (S). Consequently, the correct and incorrect peptide-MHC complex (pMHC), or S and S′, are not present at the same time during an encounter, *i.e*., they do not compete with each other for TCR binding. Second, the APC and the T cell have a finite contact duration *T*. The recognition process can only occur within this time window. Lastly, the activated states produce identical products, *i.e*., downstream signals, regardless of whether the substrate (epitope) is the correct one or not. Therefore, the cell needs a strategy to discriminate the correct and incorrect substrates based on the number of products generated within the contact time *T*.

First, assume that *T* is fixed as in previous literature [10] (we will relax this assumption later on). *Within the time window T*, the reaction diagram is represented by Eq. (4).

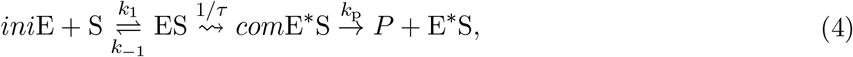

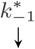Typically, TCR recognition is fairly sensitive since a few correct substrates (or epitopes) S on the APC membrane are able to activate the T cell. In the following, we will assume that each APC contains only one type of epitope, S or S′, while there are multiple TCRs that can bind to the substrate. Mathematically, the roles of substrate and TCR are equivalent. Tracking the state of the single substrate gives a similar simple stochastic process as in the DNA replication setting. This assumption allows us to reduce the number of possible states in the corresponding stochastic process.

We first consider two possible discrimination strategies, the FPT-based strategy and the product-based strategy, to quantify the output of the TCR recognition process.

### FPT-based scenario

In this strategy, reaching the activated state E*S within time *T* is interpreted as T cell activation. The output of this strategy can be represented using the FPT to E*S state (*t*_a_) as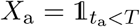. The state *X*_a_ = 1 denotes an activated T cell while *X*_a_ = 0 indicates no response by the T cell. In general, we can define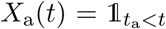, and visualize the output as a binary signal changing over time, as shown in Fig. 1(b). To justify the FPT strategy biologically, we note that upon reaching the activated state, cofactors such as CD4 and CD8 stabilize the TCR-pMHC interaction, significantly reducing the unbinding rate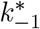. Therefore, the activated complex can then steadily produce downstream signals (products) and trigger T cell response.

### Product-based scenario

This strategy, which is implicitly analyzed in [11], estimates the pMHC-TCR “affinity” by counting the number of product or signaling molecules produced within a given time. The output of this strategy is the number of products P(*T*) at time *T*, which takes on values in ℕ, as is shown in Fig. 1(c). Consequently, in this more graded strategy, whether a T cell is activated is not described by a single, specific criterion.

Different strategies of interpreting the output requires us to define performance metrics that can compare different strategies on a common mathematical footing. We now define the performance metric that can be used to compare our two discrimination scenarios.

### Performance metrics

In the case of FPT-based discrimination, it is natural to formulate the recognition problem as a hypothesis testing problem. We denote the binary input *ξ* = 1 if S is present and *ξ* = 0 if S′ is present. The competing hypotheses are then *H*_0_ : *ξ* = 0 and *H*_*a*_ : *ξ* = 1. The cell accepts *H*_0_ if *X*_a_ = 0. There is a canonical definition of sensitivity and specificity, *i.e*., the true positive probability (TPP) and true negative probability (TNP). Given a fixed duration *T*, varying the processing time *τ* gives a family of binary classifiers (solutions to the hypothesis testing problem) corresponding to the KPR process. The receiver-operating characteristic curve (ROC) and the area under the curve (AUC) can be used to evaluate the overall performance of this family of classifiers. For a single classifier, we define the accuracy *A* as an average of specificity and sensitivity:

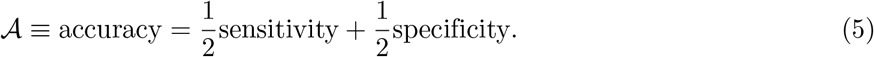

However, such metric only applies to strategies with binary output, but not the product-based strategy, as the output P(*T*) is non-binary. Kirby et al. propose a Fisher linear discriminant metric (*η*_FLD_) based on the consideration of signal-to-noise ratios [11]. *η*_FLD_ takes on values in (0, *∞*) and does not directly quantify the fidelity of transmission from the input *ξ* to the output P(*T*).

In order to compare both strategies on a common footing and represent the fidelity of discrimination directly, we introduce the mutual information ℐ between the input and output, and the associated channel capacity *C*. The mutual information between two random variables *ξ* and *X* can be defined as [23]

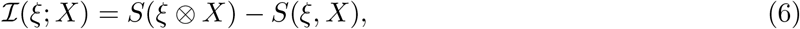

where *ξ* ⊗ *X* is the joint random variable of *ξ* and *X*, assuming that *ξ* and *X* are independent, while *S*(*ξ, X*) is the joint Shannon entropy of *ξ* and *X*. The mutual information ℐ(*ξ*; *X*) relies on the input distribution of *ξ*. Hence, one can define the channel capacity *C* as the supremum of the mutual information over all possible input distributions of *ξ* which only depends on the conditional probability distribution of *X* given *ξ*,

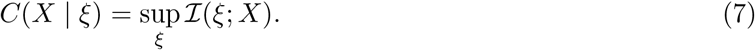

There are two advantages of using mutual information and channel capacity over the Fisher linear discriminant metric *η*_FLD_. First, the mutual information is defined for both binary variables, as in the case of the first-passage time problem, and continuous variables, as in the case of the product-based discrimination problem. Second, in the specific scenario of binary input variables (correct and incorrect substrates), the mutual information always takes values between 0 and 1 when measured in bits (log 2). A mutual information of 0 means that the distribution of input and output do not overlap while a value of 1 indicates that the distributions are identical. Consequently, the channel capacity provides a natural way to compare the product-based discrimination problem with the FPT problem and to quantify how well the system can distinguish correct substrates from incorrect ones.

In the limit in which the accuracy *𝒜 →* 1 with binary input *ξ* and output *X*, we note that the mutual information *I* is approximately *𝒜* (measured in bits) when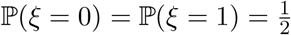 :

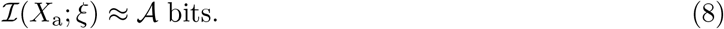

In this high accuracy limit, the channel capacity *C* is also close to the accuracy.

### Stochastic simulations

In cases where we need to rely on stochastic simulation of KPR processes, we implement the Gillespie algorithm [24] in `julia` [25]. The Gillespie algorithm tracks the state transitions of a Markov chain with exponential waiting times. In order to simulate the KPR process with a deterministic waiting time *τ*, we explicitly track all waiting times at each step and the time elapsed, updating the state by the smallest waiting time. After the state update, we re-evaluate all the waiting times and the time elapsed. The implementation is available at github.com/hsianktin/KPR.

In order to obtain the mutual information and channel capacity, we record the FPTs during the simulation, as well as the number of products produced by each simulation trajectory after a given time *T*. We simulated 10^4^ trajectories and use the empirical distributions of the FPTs and the number of products as surrogates for the true distributions. This allows us to compute the conditional probability distribution of output *X* given input *ξ*. Then, mutual information is computed using Eq. (6). The channel capacity is obtained by a numerical optimization procedure with respect to the probability of *ξ* = 1, ℙ (*ξ* =1) in the interval (0; 1).

The exact parameters used in each simulation will be specified in the corresponding figures. In general, we set *k*_*−*1_ = 1, *q*_*−*1_ = 2, indicating that the unbinding rate of an incorrect substrate is only twice as fast as that of a correct substrate.

## Results

### Long processing time improves maximal accuracy in FPT-based strategy

KPR in the DNA replication scenario relies on a comparison between the FPT to incorporate the correct nucleotide *t*_p_ and the FPT 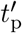 of incorporating an incorrect one, while KPR in the FPT-based TCR activation scenario requires comparison of the processing time *τ* with the cell-cell contact time *T*. As shown in Eq. (2), the error probability 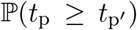 of incorporating one incorrect nucleotide scales exponentially with respect to the processing time *τ* in the *τ → ∞* limit

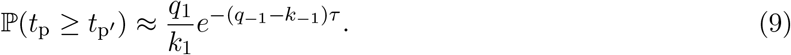

Consequently, a longer processing time *τ* can exponentially reduce the error probability, as shown in Fig. 4(a). Note that the error probability depends linearly on the binding rates *q*_1_*/k*_1_ but exponentially on the unbinding rates *q*_*−*1_, *k*_*−*1_. This different dependence indicates the nonequilibrium nature of the KPR process.

**Fig 4.**
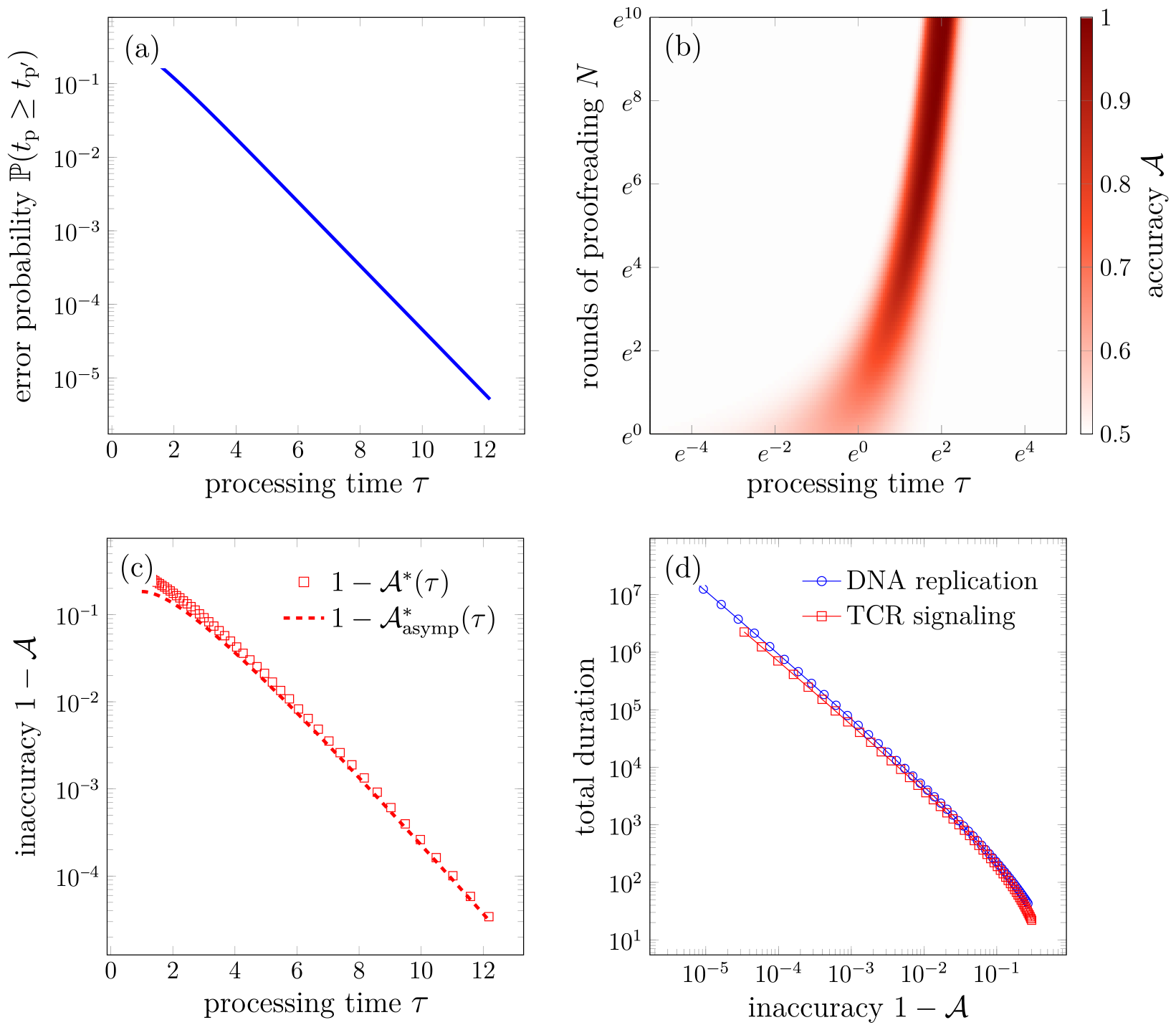
Statistics of the FPT-based strategy in the DNA replication and TCR recognition scenarios. (a) Replication error probability 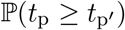 as a function of processing time *τ*, evaluated using Eq. (2). (b) Accuracy *𝒜* as a function of processing time *τ* and contact duration *T*, evaluated using Eq. (10). (c) The maximal accuracy *𝒜 \** (squares) as a function of processing time *τ*, evaluated using Eq. (12). The asymptotic behavior of *𝒜 \** in the *τ → ∞* limit is shown by the dashed curve, evaluated using Eq. (13). (d) Total duration (MFPT 𝔼[*t*]) of the DNA replication process as a function of inaccuracy 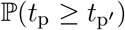 is evaluated using Eq. (3) and is shown in blue. The optimal contact duration *T* * in the TCR recognition scenario as a function of inaccuracy 1 *− 𝒜 \** (red) is evaluated using Eq. (12) and the definition *N* = *k*_1_*T*. In (a-d), we set *k*_1_ = *q*_1_ = 0.1, *k*_*−*1_ = 1, and *q*_*−*1_ = 2.

For convenience, in FPT-based scenario, we assume parameters 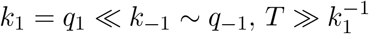 and let *N* = *Tk*_1_, which allows us to treat the recognition process as *N* cycles of a one-shot process. For each shot, an initial complex ES either unbinds with a probability of 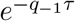 or 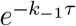, or activates. Then, *N* independent cycles of the process yields the geometric success probability 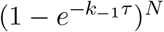.

Under the assumptions introduced previously, we can evaluate the accuracy *𝒜* defined by Eq. (5) as

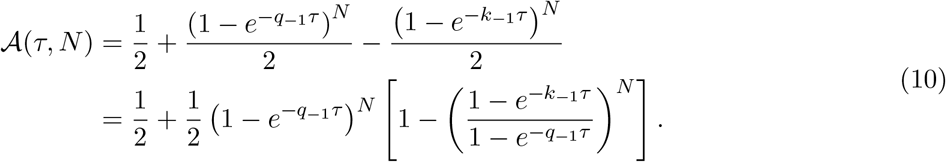

The accuracy depends on both the number of binding events *N* (hence *T*) and processing time *τ*, as illustrated in Fig. 4(b). For fixed contact duration *T*, the accuracy first increases with *τ*, then followed by a decrease. In the long processing time limit *τ → ∞*, the T cell does not respond to any signal, corresponding to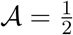.

For fixed processing time *τ*, there is a maximal accuracy 𝒜 *\**(*τ*) = sup_*N*_ 𝒜 (*τ, N*) and a corresponding *N* *(*τ*) such that 𝒜 (*τ, N* *) = 𝒜 *\**(*τ*). A straightforward calculation yields

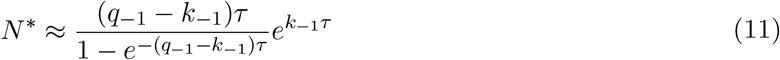

and

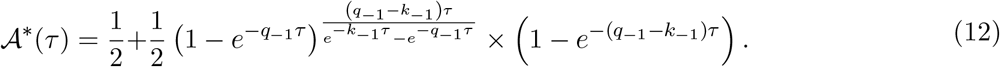

The asymptotic behavior of 𝒜 *\** in the *τ → ∞* limit is

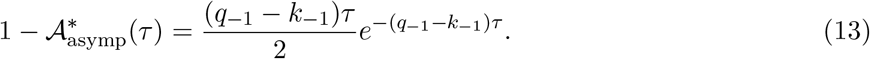

As demonstrated by Eq. (9), Eq. (13), and Fig. 4(a,c), for both DNA replication and TCR recognition scenarios, the *inaccuracy* or error probability scales with 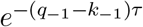. The improvement in the accuracy comes at a cost of the increased total times spent in the proofreading process. The mean time spent in the DNA replication process is given by Eq. (3). The optimal contact duration *T* * required for a specific *τ* is given by 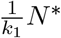, with *N* * given by Eq. (12). In both cases, there is a common 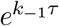 factor. Thus, for both scenarios, there is a trade-off between the accuracy and the total time spent in the proofreading process, as is shown in Fig. 4(d).

In the following sections, we will focus primarily on the TCR recognition scenario.

### Channel capacity and Fisher linear discriminant agree qualitatively

We first show that the channel capacity metric and the Fisher linear discriminant metric show qualitatively similar behavior. To obtain statistically accurate results, we simulate the TCR recognition process using the Gillespie algorithm and evaluate the channel capacity and Fisher linear discriminant metric from 10^4^ realizations. The results shown in Fig. S2 and Fig. S3 indicate that both quantities increase with respect to cell-cell contact time *T* and are maximal at waiting time *τ ∼* 0.6*/k*_*−*1_, regardless of *T*. We now establish the channel capacity as an appropriate metric for comparing different discrimination strategies.

### Invariant optimal processing time of product-based strategy for different contact times

We now compare the channel capacity of FPT-based discrimination to that associated with product-based discrimination. In Fig. 5 we plot the channel capacity between the input *ξ* and the outputs 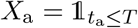 (in the FPT-based discrimination) and P(*T*) (in the product-based discrimination) as a function of cell contact time *T* when *τ* = 3 is fixed.

**Fig 5.**
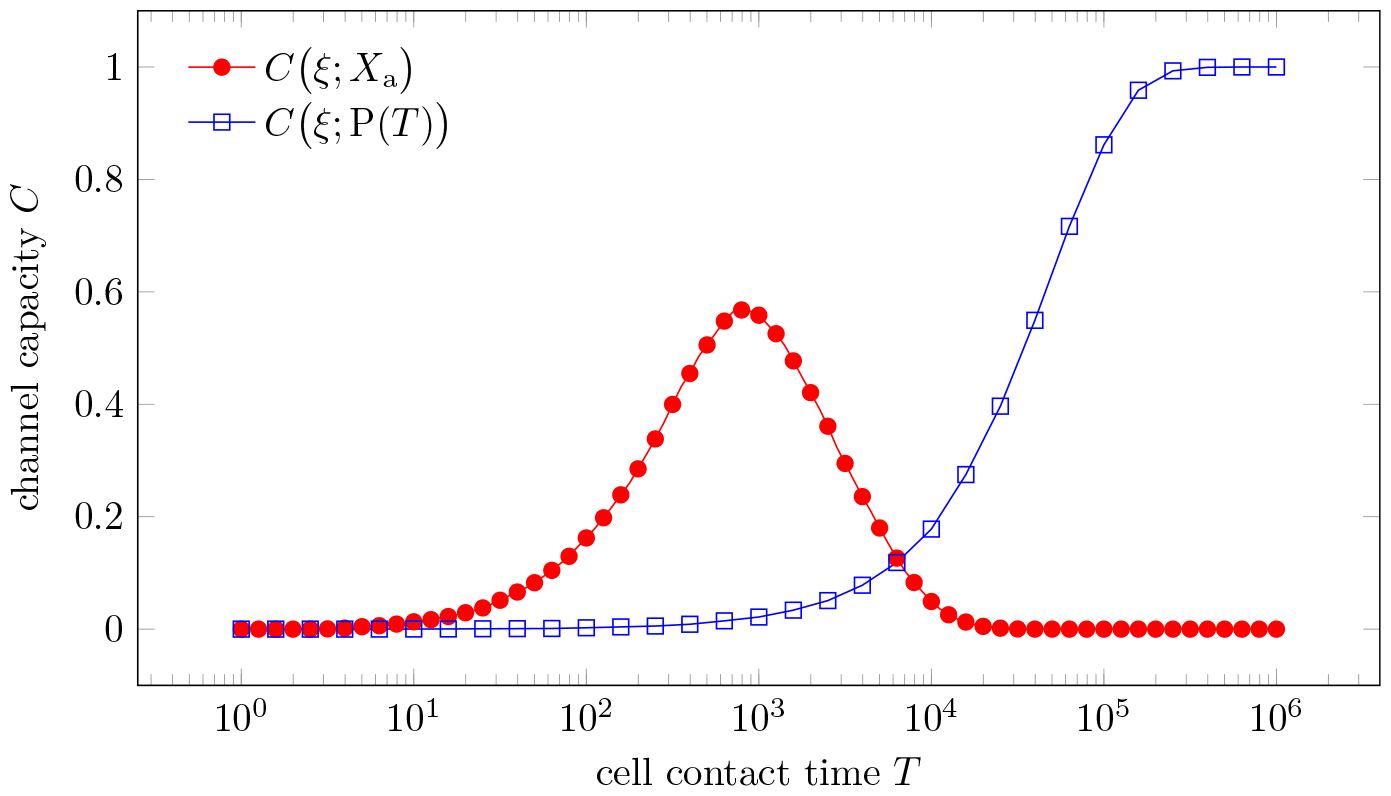
The channel capacity as a function of cell-cell contact time *T* for first-passage-time-based (FPT-based) signaling and product-based signaling. The channel capacity is evaluated between the input *ξ* indicating correct (1) or incorrect (0) substrate and the output *X*_a_ or P(*T*). We assumed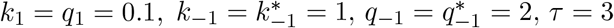, and *k*_p_ = 0.01 for a slow product formation rate.

In Fig. 6, we plot the channel capacity as a function of processing time *τ* for various contact times *T*. As in Fig. 5, we channel capacities associated with both FPT-based and product-based discrimination. The channel capacity of product concentration increases monotonically with respect to cell contact time *T* while exhibiting a peak at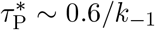, as is shown previously. By contrast, the channel capacity under FPT-based discrimination has an optimal processing time 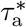 that increases with respect to cell contact time *T*. The optimal contact time 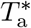 also increases with respect to the processing time *τ*. The channel capacity-optimizing contact times 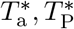, and processing times 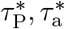 shown in Fig. S1(a,b).

**Fig 6.**
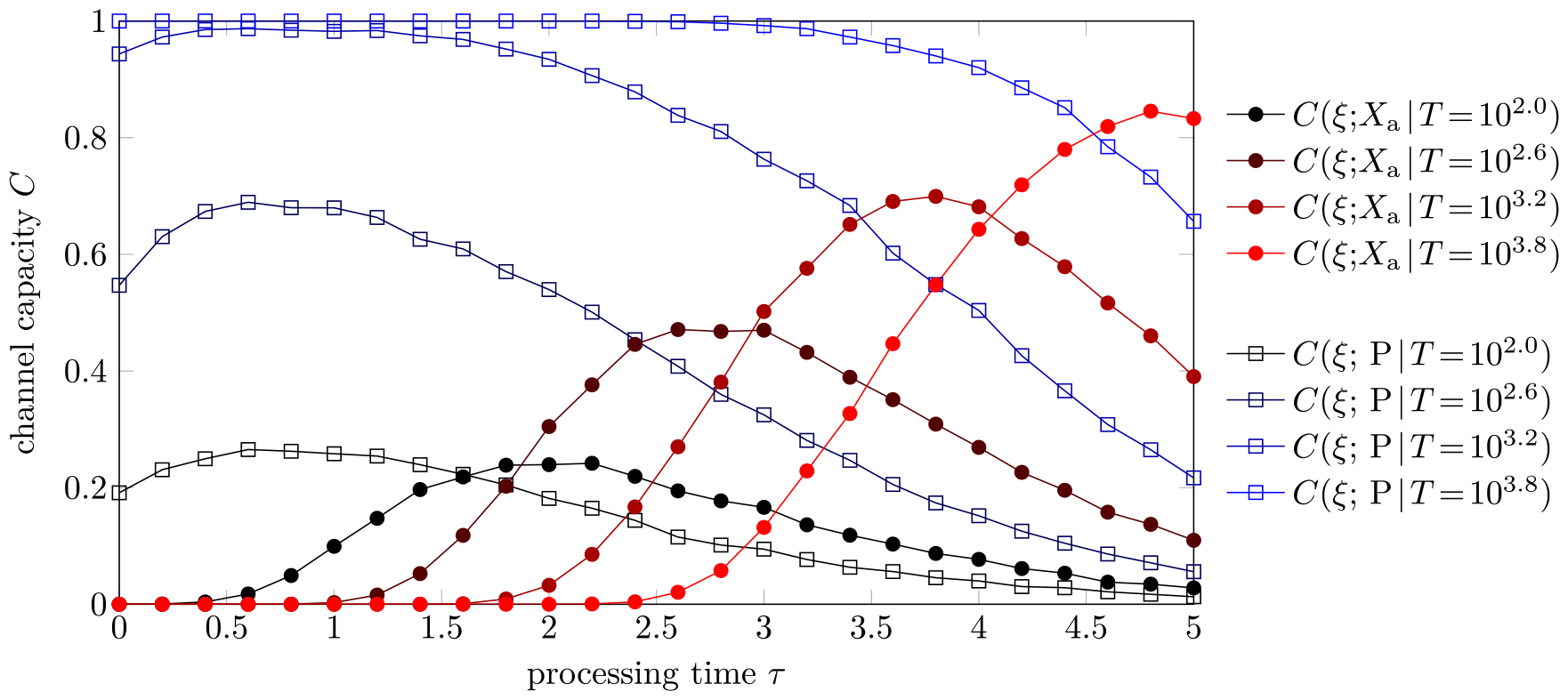
The channel capacities of product concentration (blue squares) and first activation times (red dots) as a function of processing time *τ* for various cell contact times *T*. We evaluate the channel capacity using stochastic simulations (Gillespie algorithm) of the model in Eq. (4) with parameters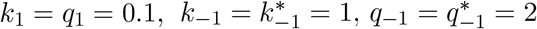, and *k*_p_ = 1. The production rate was set higher for easier simulation of the product concentration.

There are two noteworthy observations from Fig. 6. First, in the product-based scenario, the channel capacity for *τ* = 0 (no kinetic proofreading limit) does not differ significantly from the optimal channel capacity at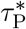. This observation together with the *T* -independent optimal processing time 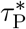 reflects the conclusion by Kirby et al. that KPR is ineffective due to noise [11]. Second, under the same total contact duration *T*, the maximal channel capacity of the product-based strategy is higher than that of the FPT-based strategy provided that the production rate *k*_p_ is sufficiently large. These two observations suggest that the product-based discrimination is a superior strategy compared to the FPT-based one. In order to provide mechanistic insight into the difference between these two strategies, we now introduce a method to analytically characterize the product-based strategy.

### Decomposition of the product-based strategy

We propose decomposing the product-based strategy into a series of first-passage-time-based strategies using the FPT of the number of products P to different thresholds P_th_. Let *t*_*k*_ represent the first time the product P(*t*) exceeds the threshold *k. X*_th_ = 1 (triggering immune response) if *t*_P_th *≤ T* and *X*_th_ = 0 otherwise. Being a FPT-based strategy, *C*(*ξ*; *X*_th_) has a similar dependence on *T* and *τ* to that of *C*(*ξ*; *X*_a_), the channel capacity in FPT-discrimination we discussed earlier. In order to illustrate this point, we perform Gillespie simulations of the model in Eq. (4) with the same parameters as in Fig. 6. The results are shown in Fig. 7 and Fig. 8.

**Fig 7.**
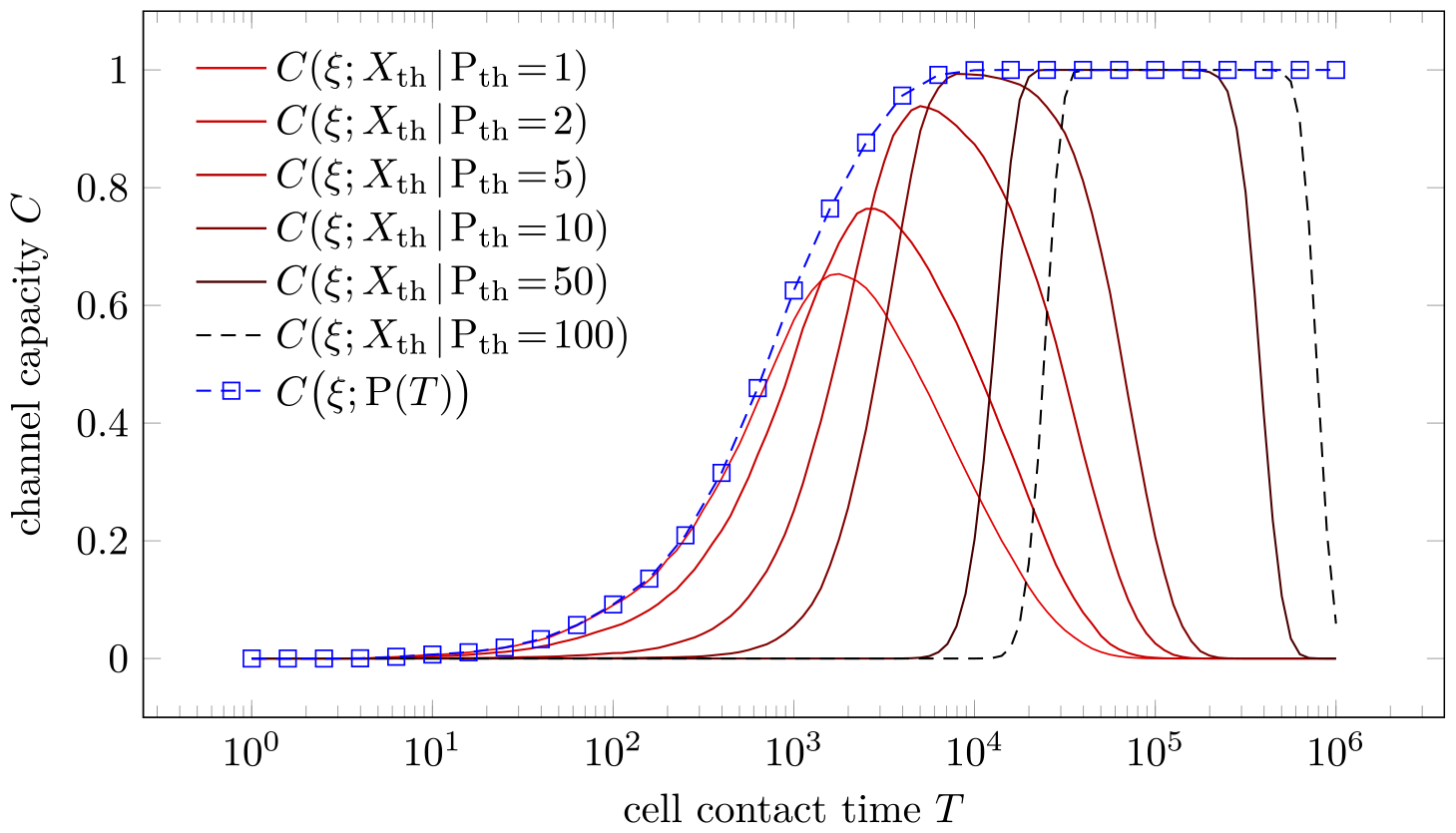
The channel capacity between the input *ξ* and the output P(*T*) or *X*_th_ as a function of cell-cell contact time *T*. Here, 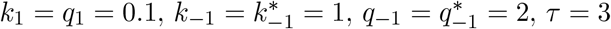, and *k*_p_ = 1.

**Fig 8.**
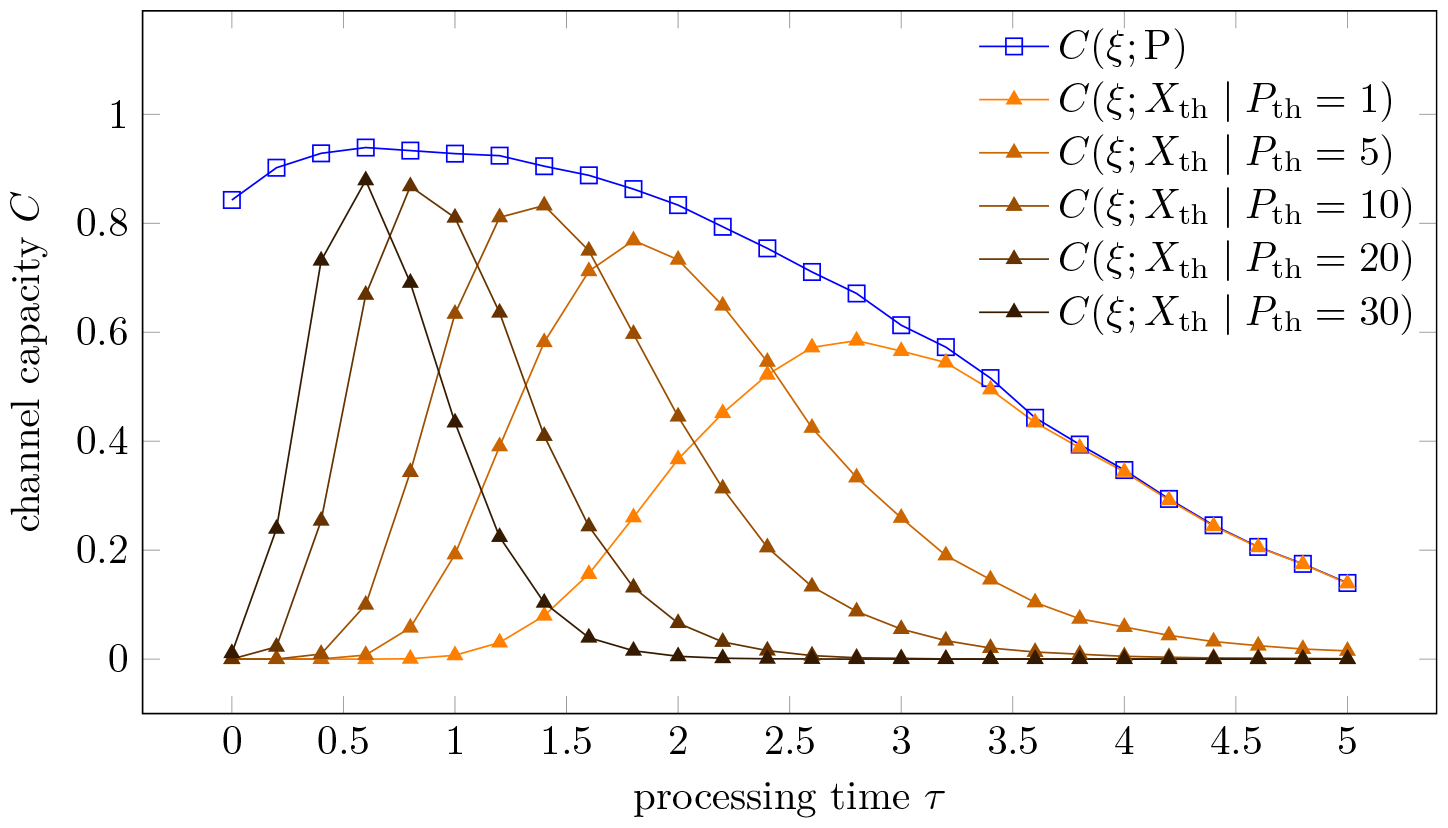
The channel capacity between the input *ξ* and the output *X*_a_, P(*T*), or *X*_th_ as a function of processing time *τ*. We assumed 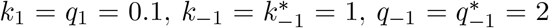, *T* = 1000, and *k*_p_ = 1. 10,000 independent Gillespie simulations are conducted for each *τ*.

Since *X*_th_ is derived from P(*T*), *C*(*ξ*; P(*T*)) serves as an upper bound of *C*(*ξ*; *X*_th_) for various thresholds *P*_th_, as is illustrated in Fig. 7 and Fig. 8. No single threshold can reach the information upper bound *C*(*ξ*; P(*T*)) at any *T*. A small threshold P_th_ can approach *C*(*ξ*; P(*T*)) for small *T* and large *τ*, while a large threshold P_th_ can approach the information upper bound *C*(*ξ*; P(*T*)) for large *T* and small *τ*. We thus introduce the approximation

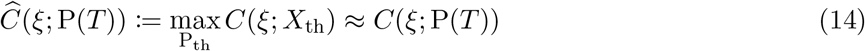

for *C*(*ξ*; P(*T*)).

### Mathematical analysis of the effects *τ* in the product-counting strategy

Having established the approximation to the channel capacity of the product-based strategy in Eq. (14), we further analyze its dependence on the processing time *τ* under a fixed cell-cell contact time *T*. We use the equivalence between channel capacity and accuracy in the high accuracy limit, as shown in Eq. (8) to evaluate *C*(*ξ*; P(*T*)). We first introduce a Gaussian distribution approximation to the original distribution of P(*T*) with matched mean and variance

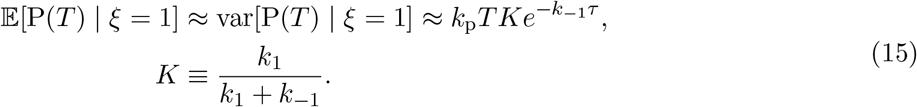

The *ξ* = 0 case is similar. The above steady-state approximation is justified in Appendix A1.

The condition *t*_P_th *< T* is equivalent to P(*T*) *≥* P_th_. Approximating the distribution of P(*T*) by a normal distribution with mean 𝔼 [P(*T*)] and variance var [P(*T*)], we can estimate the conditional probabilities of 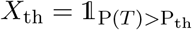 given ξ by the integral

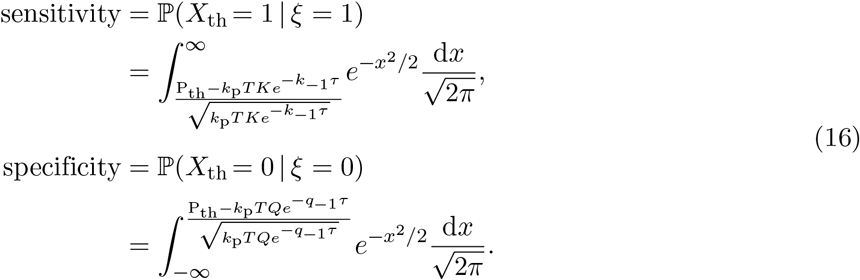

Under the assumption that 𝔼[P(*T*) | *ξ* = 1] *>* P_th_ and 𝔼[P(*T*) | *ξ* = 0] *<* P_th_, we can rewrite Eq. (16) as

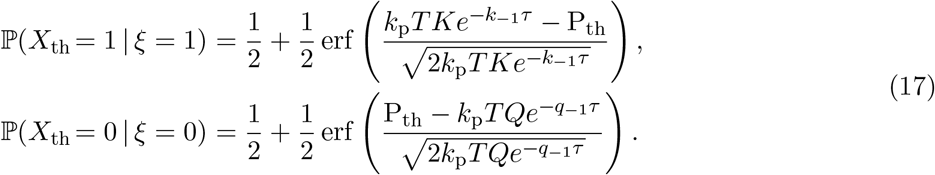

In the high accuracy limit, the mutual information *C*(*ξ*; *X*_th_) between binary uniform input and binary output is approximated by the accuracy 𝒜 as in Eq. (8). We can then obtain the optimal threshold 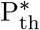 that maximizes the accuracy 𝒜 _th_ = ℙ(*X*_th_ = 1 | *ξ* = 1) + ℙ(*X*_th_ = 0 | *ξ* = 0) for given *T* and *τ* to be

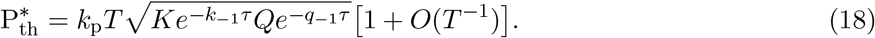

The exact optimal threshold 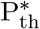 can be analytically solved, but we keep the above asymptotic form for simplicity when *T → ∞*. The maximal accuracy 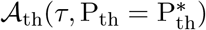 is then used to approximate the maximal channel capacity 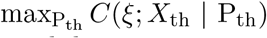, which is an approximation of the channel capacity *C*(*ξ*; P(*T*)) of the product-based discrimination.

Substituting the zero-th order term of Eq. (18) into Eq. (17), we find

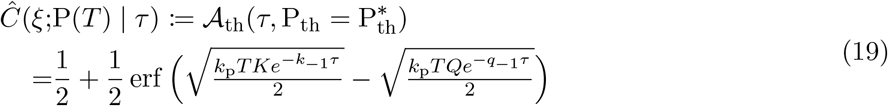

In Fig. 9, note that Eq. (19) generally matches the simulation results well. Slightly higher values of Eq. (19) arise from the Gaussian approximation used to map discrete output to continuous output.

**Fig 9.**
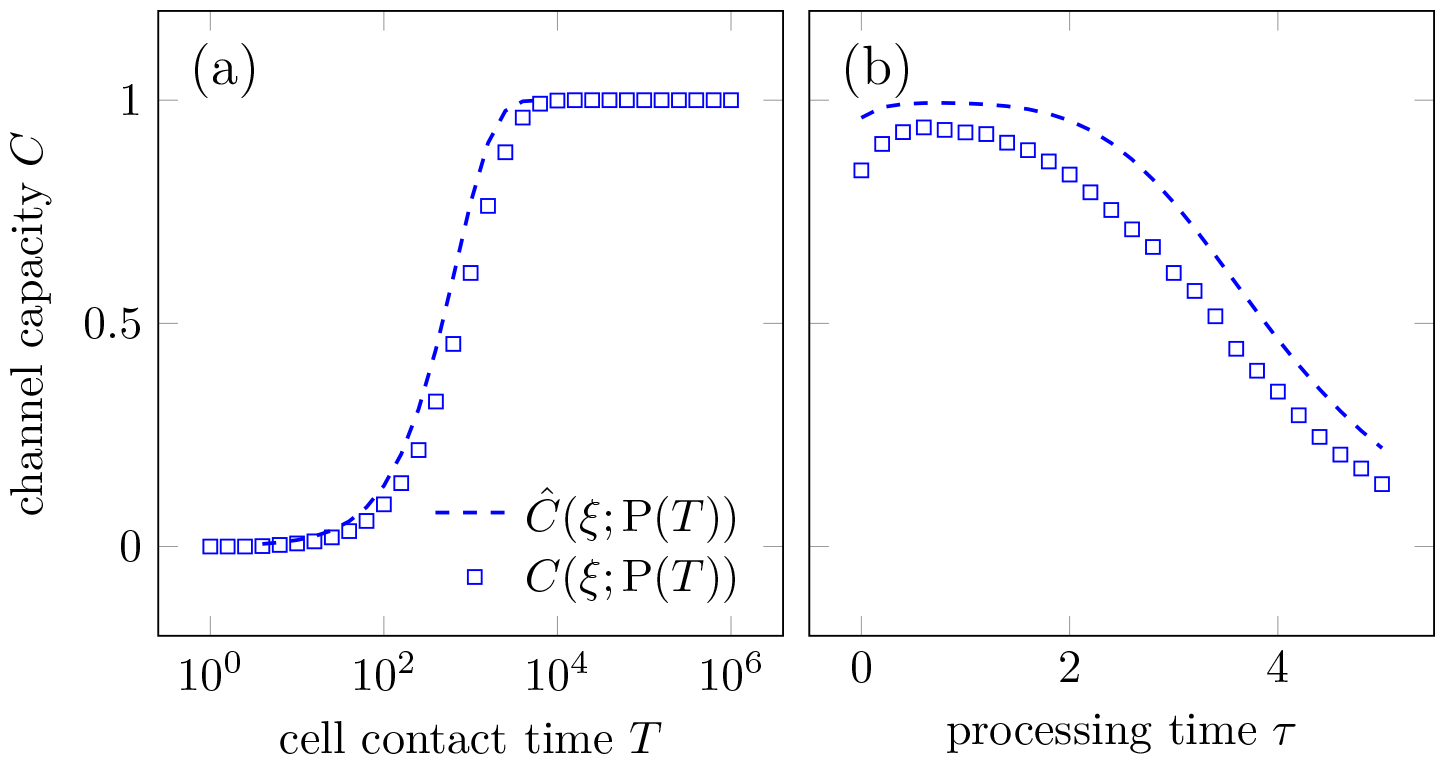
Comparison between the simulated channel capacity *C* (*ξ*; P(*T*)) and the corresponding estimate using Eq. (19). (a) *C*(*ξ*; P(*T*)) as a function of cell-cell contact time *T* ; (b) *C* (*ξ*; P(*T*)) as a function of processing time *τ*. Here, we took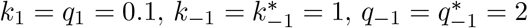, and *k*_p_ = 1. *τ* = 3 in (a) and *T* = 1000 in (b).

The choice of optimal processing time 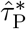 that maximizes the accuracy in Eq. (19) is independent of *T* and is given by

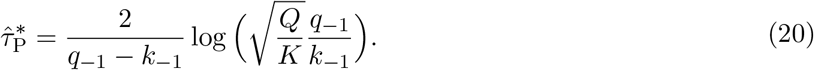

Note that the optimal processing time 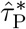 is obtained by taking 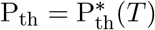 and is independent of the cell-cell contact time *T*. Under the parameter settings where *k*_1_ = *q*_1_ = 0.1, *k*_*−*1_ = 1, and *q*_*−*1_ = 2, we find that 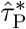 is approximately 0.6*/k*_*−*1_, which is consistent with the optimal processing time 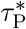 obtained from the simulation results, as illustrated in Fig. S1.

### The dynamical threshold and random cell-cell contact time

In the previous sections, we have assumed that the cell-cell contact time *T* is deterministic. In reality, the cell-cell contact time *T* is random and vary from cell to cell [26–28]. To evaluate the effects of a random cell-cell contact time, we consider a simple model where the cell-cell contact time *T* is uniformly distributed in the interval [0, *T*_max_], where *T*_max_ is the maximal cell-cell contact time.

In the previous section, we conclude that the *T* -independent optimal processing time 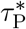 is a result of fixing the threshold P_th_ to 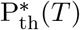 given by Eq. (18). In the case of a random cell-cell contact time *T*, choosing a universally optimal threshold 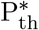 is difficult. We can, however, choose a dynamical threshold 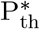 (*t*) that increases with the time *t* passed since the initial contact to maximize the channel capacity, where 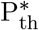 is still given by Eq. (18), with *T* replaced by *t*. A comparison of a dynamical threshold 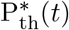 and a static threshold P_th_ is shown in Fig. 10(a).

**Fig 10.**
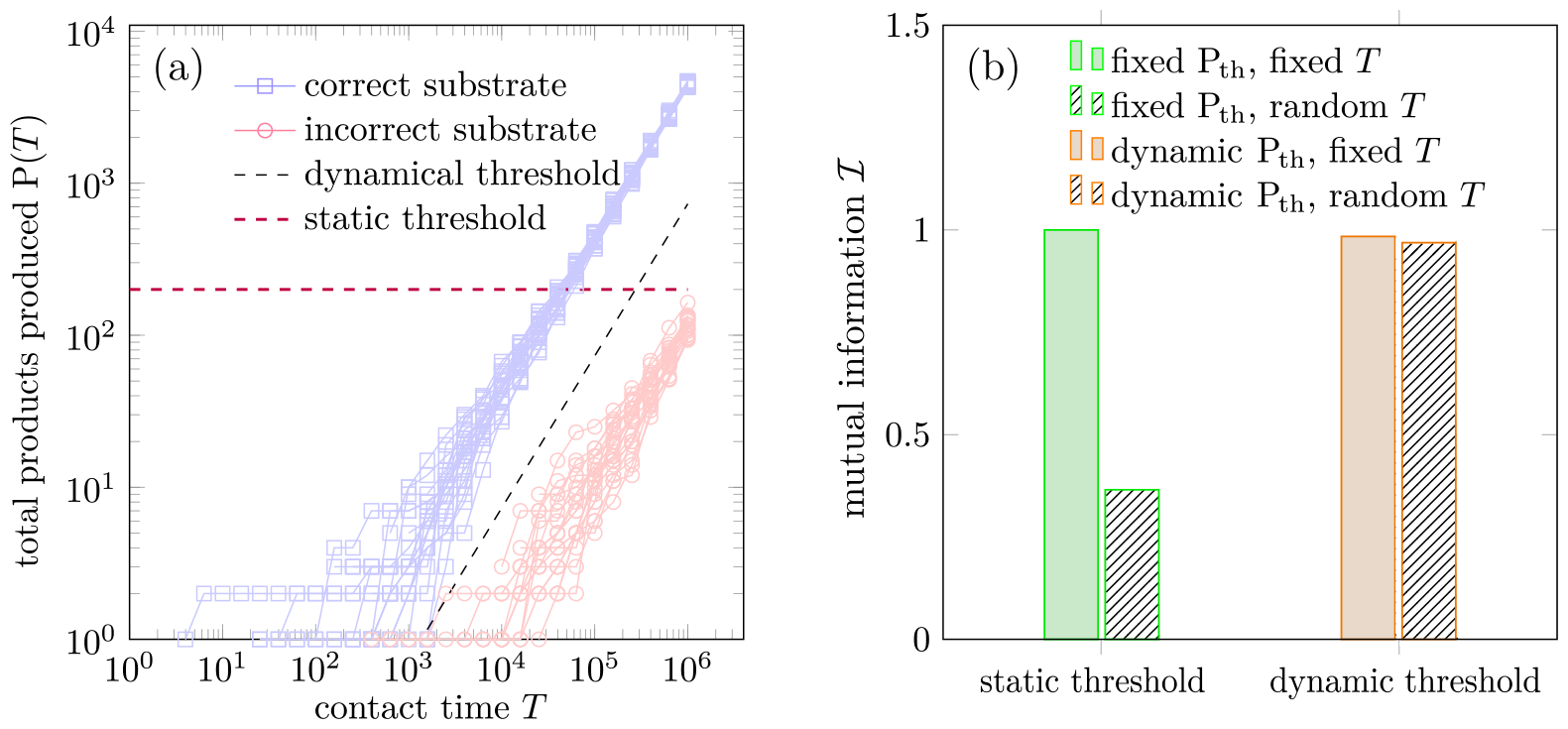
The dynamical threshold and random contact time. (a) Illustration of the dynamical threshold 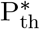 as a function of the time *t* since initial contact. The blue trajectories represent the number of products P with correct substrates. The red trajectories represent that of incorrect substrates. (b) A dynamical-threshold-based discrimination strategy maintains a high channel capacity when the total contact time *T* is uniformly distributed between 0 and *T*_max_. Filled bars represent the mutual information between input *ξ* and output *X*_th_ with a fixed contact time *T* and patterned bars represent the mutual information with a uniformly distributed contact time *T* between 0 and *T*_max_. The green bars indicate the maximal mutual information over all possible contact times *T ≤ T*_max_ and all possible static thresholds P_th_. The input *ξ* is assumed to be uniformly distributed on {0, 1}. We assumed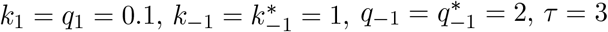, and *k*_p_ = 1. *T*_max_ is set to 10^6^. We additionally mandate that when the dynamical threshold 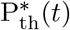 is smaller than 10 products, no response is initiated to filter out the initial noise.

The numerical results of the maximal mutual information between the input and output *X*_th_ under different contact times *T* and different thresholds P_th_ are shown in Fig. 10(b). In the case of a fixed contact time *T*, the maximal mutual information of both static and dynamical thresholds is close to 1, indicating perfect discrimination. In the case of a uniformly distributed contact time *T* between 0 and *T*_max_, the maximal mutual information of the dynamical threshold is close to 1, while that of the static threshold is close to 0.4, indicating poor discrimination.

The experiments in Fig. 10 suggest that the dynamical threshold 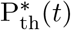 maintains a high channel capacity when the total contact time *T* is random.

### A nested single-binding KPR scheme

We have provided a mechanistic explanation for the different qualitative behaviors of the channel capacity between first-passage-time-based discrimination and product-based discrimination, in particular, why the optimal processing time 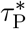 is independent of the cell-cell contact time *T* in the product-based discrimination.

It is also of interest to understand the high channel capacity in the product-based discrimination problem that arises even in the *τ →* 0 limit, as shown in the blue curve of *C*(*ξ*; P) in Fig. 8. To provide simple mechanistic intuition, note that production and accumulation of products can be comparable to a sequence of phosphorylation events in the traditional KPR scheme. The cell-cell contact time *T* sets the rate of the competing unbinding event:

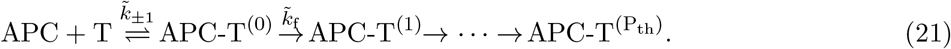

Here, ATC represent the antigen-presenting cell (APC) and the T cell is denoted by T. The superscript (*i*) represents the number of products generated by the T cell. The T cell initiates responses when the number of products exceeds a threshold P_th_. The overall scheme is comparable to the traditional KPR scheme shown in Fig. 2(a). The difference between the traditional KPR scheme and the product-based strategy shown above is that the latter assumes a different phosphorylation rate 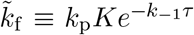 or 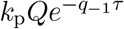 for correct and incorrect substrates, while the unbinding event has a common timescale *T* set by the cell-cell contact time.

We have assumed that the contact time *T* is approximately constant given that unbinding of the TCR and the APC requires multiple steps involving collective effects of adhesion membrane proteins. Also note that the activation time of the TCR signaling process, given by the FPT 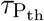 to generate a number of products P beyond the threshold P_th_ is also approximately a constant when P_th_ is sufficiently large. Consequently, the competition between two processes with almost fixed timescales allows highly informative outputs compared to the traditional KPR scheme and Michaelis-Menten kinetics, as indicated in Eq. (22), where 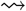 and 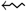 represent a deterministic waiting time *τ*.

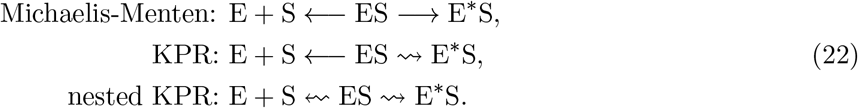

To further illustrate the role of a deterministic waiting time in the accuracy of the above process, we compare it with an exponential waiting time, *i.e*., standard Michaelis-Menten (MM) kinetics in Appendix A2. As for a comparison between the traditional KPR scheme and the nested KPR scheme, we note that in the limit of deterministic waiting time, the nested KPR can achieve exact discrimination.

## Discussion

In this paper, we have considered kinetic proofreading schemes for two classes of biological processes, DNA replication process and T cell signaling. Overall, our analysis indicates that how the output of a kinetic proofreading process is used to make a decision is crucial to the performance of KPR. We have shown that in the case of DNA replication, the specificity of the process is always exponentially dependent on the processing time *τ* and increasing specificity comes at the cost of replication speed.

In the case of TCR signaling, the trade-off between specificity and sensitivity can be mitigated by increasing the number of allowed failed attempts *N* which is proportional to the cell-cell contact time *T*. The overall accuracy 𝒜 of the signaling process is still exponentially dependent on the processing time *τ*, as illustrated by Eq. (13). For longer processing time *τ*, the higher accuracy can only be achieved at the cost of the signaling speed by exponentially increasing the cell-cell contact time *T*, or equivalently, the number of allowed failed attempts *N* to the optimal value *N* * indicated by Eq. (11).

A variant of the FPT-based strategy, extracting the extreme FPT in the presence of multiple substrates (antigens), has also been proposed [10]. The main goal in [10] is to determine sensitivity and specificity of TCR recognition of foreign antigens when also exposed to a sea of self-antigens. The self-antigens in [10] are assumed to bind much more weakly to the TCR than the foreign antigens. While we are primarily interested in discrimination between correct and incorrect substrates with similar binding affinities. This is a typical situation as T cells need to identify cancer cells that present mutated self antigens on their surface. In this case, the affinity of the corresponding TCR to the self antigen is expected to be similar to the affinity to the foreign antigen. The densities of the self and foreign antigens are also expected to be similarly low.

The amount of product produced can also be used to distinguish correct and incorrect substrates. To compare the performance of product-based discrimination to that of FPT-based discrimination, we introduced a channel capacity between the input *ξ* ∈ {0, 1} (incorrect, correct substrate) and the output *X*, which can be either 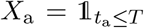 or the number of products generated P(*T*). We established the connection between the channel capacity and the accuracy of the discrimination problem in the high accuracy limit in Eq. (8). Thus, the dependence of the first-passage-time-based discrimination on the processing time and the cell-cell contact time can be shown to have a single peak, corresponding to the optimal processing time 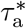 and the optimal cell-cell contact time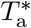, respectively, as illustrated in Figs. 5–6.

By contrast, the product-counting discrimination problem has a distinct monotonic increase of the channel capacity with respect to the cell-cell contact time *T* and the optimal processing time 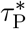 is independent of *T*. We find analytic approximations of the channel capacity by decomposing the product-based discrimination problem into a series of first-passage-time-based discrimination problems with different thresholds. This observation allows us to analytically approximate 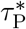 by Eq. (20). The approximation in turn reveals that the peculiar dependence of the channel capacity on the processing time and the cell-cell contact time arises from choosing an optimal threshold 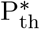 for distinguishing correct substrates from incorrect ones.

Our TCR signaling model is a simplified scenario that assumes a deterministic cell-cell contact time *T* and a deterministic processing time *τ*. However, we conducted additional simulations that relaxed the deterministic processing time assumption by explicit modeling of the irreversible phosphorylation process in Appendix A3. The results show qualitatively similar behavior as the simplified model. Additionally, we fixed the mean processing time and varied the number of processing steps *m* to explore the effect of the number of processing steps in KPR on the channel capacity. The results suggest that larger *m* increases the channel capacity of the FPT-based discrimination scheme, while the channel capacity of the product-based discrimination remains effectively unchanged, as shown in Fig. S4.

A random cell-cell contact time *T* may also impair the performance of the T cell in distinguishing correct ligands from incorrect ones. The dynamical threshold P_th_ may rescue this impaired performance by allowing the T cell to effectively adjust the threshold in time. This rescue is illustrated in Fig. 10. However, implementing a dynamic threshold requires the T-cell to keep track of the duration since the initial contact with the APC to adjust the threshold P_th_ accordingly. The duration may be tracked by another series of similar mechanical or biochemical reactions on the membrane-membrane interface that are triggered by the membrane-membrane contact, as discussed in [29]. Experimentally, the presence of such a dynamical threshold can be detected by simultaneously measuring the number of products P(*T*), the total contact time *T* until full activation, and other markers indicating whether the T-cell is activated or not.

Alternatives to the dynamical threshold strategy have been introduced in the literature, including adaptive KPR models [12–16], force-dependent signaling including catch bonds [30, 31], and KPR through spatial gradient [32]. Our analysis approach can also be applied to these different contexts to provide better analytical understanding of the interplay between speed and accuracy.

The main observation that motivates the introduction of different strategies for discrimination is that TCR signaling is not an isolated process. Rather, signaling processes are an integral part of the cellular reaction network. It would be interesting to explore how the information is transferred from the TCR signaling process to the downstream reaction networks. Such investigations may provide insights into a long-standing but often overlooked question: what type of information is transferred from signaling process to the downstream pathways? Our work assumes that information is primarily transmitted via a binary signal that decides whether a T cell response is triggered, while Kirby et al. [11] assumed that the information is a continuous signal that reflects the strength of binding affinity between ligands and receptors.

On a theoretical level, kinetic signaling schemes represent stochastic, biological implementations of the classical Maxwell’s demon [33], where the receptor is the demon that measures the affinity of the ligand to the receptor and sorts the ligands accordingly. The canonical Maxwell demon needs memory to measure both the position and time of a particle. In the case of a stochastic demon and the measurement of binding affinity, a memoryless exponential processing time seems to be able to provide a nonzero channel capacity to distinguish the correct and incorrect ligands. However, additional memory as provided by the nonequilibrium kinetic proofreading process (and non-exponentially distributed waiting time *τ*) enhances the channel capacity significantly. While previous literature has explored the idea of a Maxwell’s demon [34, 35] and energy-accuracy bounds in generalized KPR processes [36], the quantitative interplay between energy cost, memory, and information processing still lacks a suitable language and awaits future elucidation. In particular, we have not considered energy cost which may influence the preference in cells for different strategies of discriminating correct substrates from incorrect substrates.

## Supporting information

**S1 Appendix. Mathematical appendices and additional figures. Appendix A1** master equation for the simplified KPR model. **Appendix A2**: analysis of information transmitted by KPR and Michaelis-Menten schemes. **Appendix A3**: multistep binding model. **Appendix A4**: additional figures.

## Acknowledgments

The authors thank A. Zilman and X. Guo for insightful discussions on KPR and the manuscript.

## S1 Supplementary Information

### A1 Master equation for the stochastic KPR model

Here, we consider the master equation associated with the multi-round proofreading model described in the TCR setting of the main text and derive numerical methods to evaluate the mean and variance of the product P(*T*) produced up to time *T*.

We use ℙ(*n*, E + S; *t*) to denote the probability at time *t* that the system in the E + S state and that exactly *n* products exist. Similarly, we use ℙ(*n*, ES; *t*) and ℙ(*n*, E*S; *t*) to denote the probability of *n* products and the system in the ES and E*S states, respectively. However, since our model involves a deterministic processing time *τ*, we introduce the probability density function *ρ*(*n, a*; *t*), where *a* indicates the age of the complex since its formation. ℙ(*n*, ES; *t*) and ℙ(*n*, E*S; *t*) are then given by

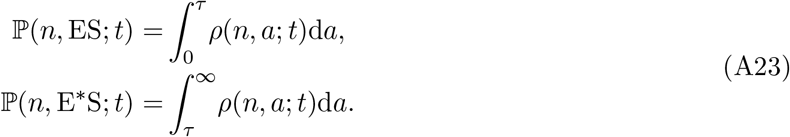

The off rates can be considered as a death rate of the age-structured complexes ES and E*S, while the on rate *k*_1_ times the probability of the E + S state is a birth rate of ES. Consequently, an age-structured master equation can be written as

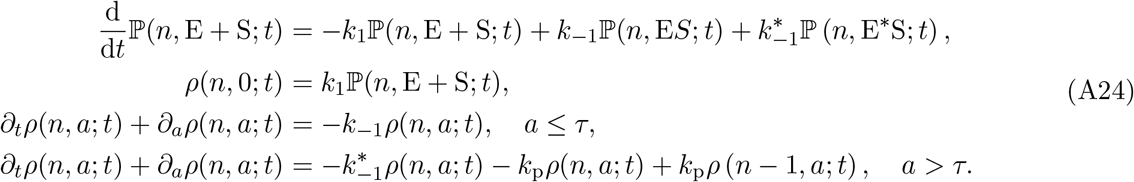

The average of the number of products P at time *T* is defined by

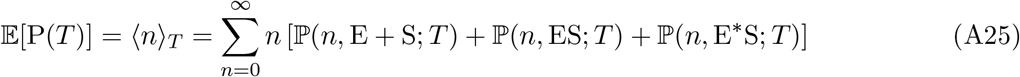

and satisfies the ODE

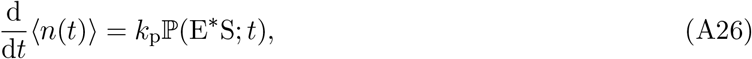

where 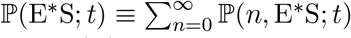 is the marginal probability of the E*S state. Similarly, to find the variance of P(*T*), we first find the dynamics of *⟨n*^2^(*t*) *⟩* as follows

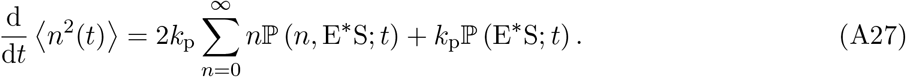

We denote 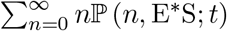 by ⟨*n*(*t*)|E*S⟩. Similarly, we have 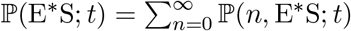 and the like. We develop a system of equations to solve for ℙ(E*S; *t*):

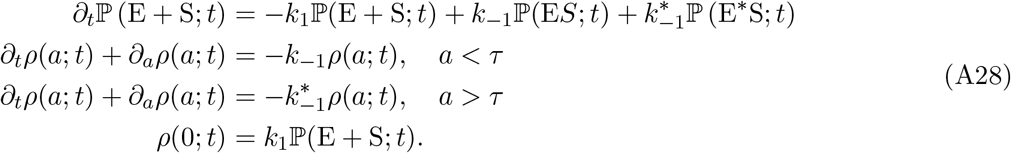

Since we are interested in finding ℙ(E*S; *t*) alone, we can integrate Eqs. (A28) over age *a* to find the reduced system of equations

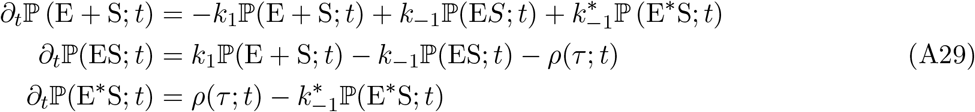

Eq. (A29) is not closed because of the density term *ρ*(*τ* ; *t*), but we can use a mean-field approximation by assuming that the internal relaxation of *ρ*(*a*; *t*) is very fast. Then, we have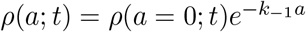, allowing us to approximate

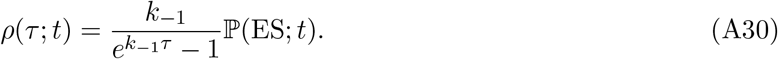

Substituting Eq. (A30) into Eq. (A29), we obtain a closed system of equations for ℙ(E*S; *t*) and ℙ(ES; *t*). The approximation is valid in the sense that the steady state of the system is consistent with the steady state of full system described by Eq. (A28). In particular,

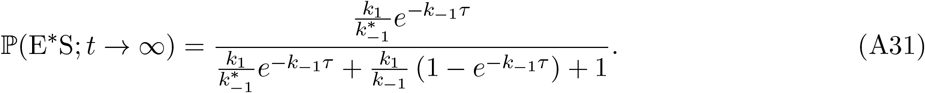

Similarly, we develop a system of equations as follows to solve for ⟨*n*(*t*)|E*S⟩ using the same approximation in Eq. (A30):

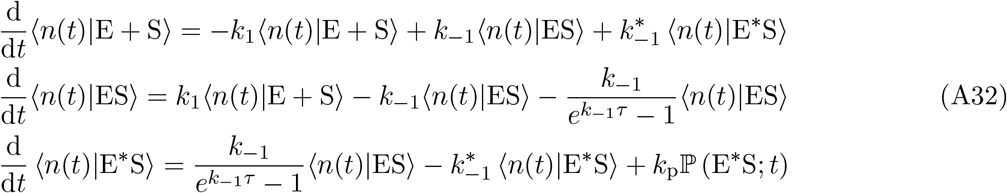

The initial conditions are given by ⟨*n*|*Y* ⟩_0_ = 0 for *Y* = E +S, ES, E*S. The analytic solutions to Eq. (A32) are not tractable but formal solutions can be obtained by Laplace transform. Uisng the approximation ℙ(E*S; *t*) *≡* ℙ(E*S; *t → ∞*) given in Eq. (A31), the Laplace transform of Eqs. (A32) is given by

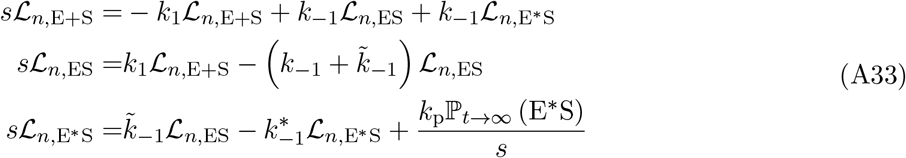

where 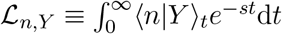 and 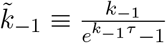. The solution can be expressed as

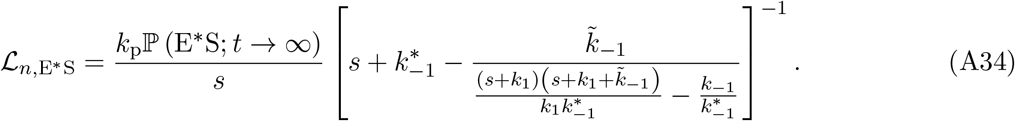

#### A1.1 *k*_p_ *→* 0 **limit**

Taking the *k*_p_ *→* 0 limit, we find

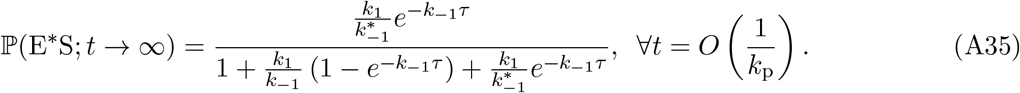

Equation (A26) gives

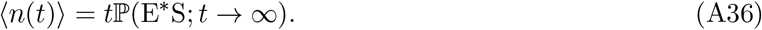

Additionally, separation of time scales allows us to approximate the solution to Eq. (A32) with

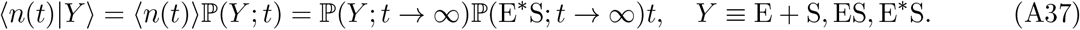

In particular, Equation (A27) becomes

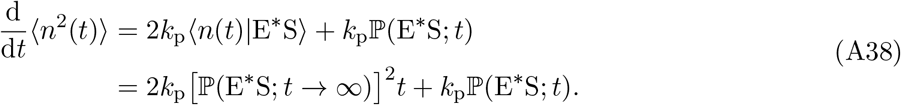

Since ⟨*n*^2^(*t* = 0) ⟩ = 0, we can integrate the above equation to find

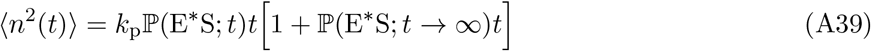

In other words, the variance of P at time *t* is given by

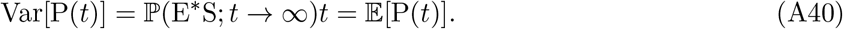

One can further verify that the distribution of P(*t*) should be Poisson in the limit *k*_p_ *→* 0.

#### A1.2 Limit of discrete processes

As we have discussed, our deterministic processing time *τ* can be considered as a limit of a discrete *m*-step KPR process with 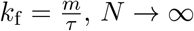. provided expressions for the mean and variance of the product P at time *T* when 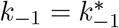:

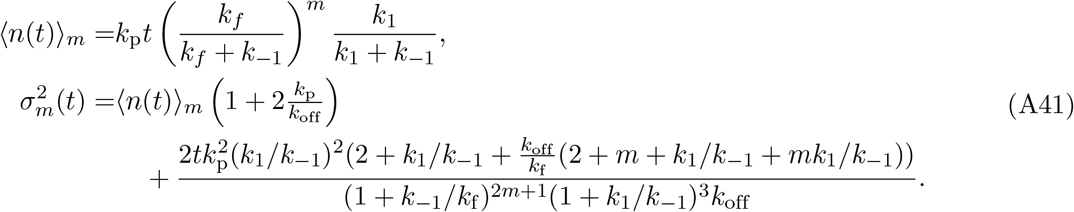

In the limit of *m → ∞*, we have

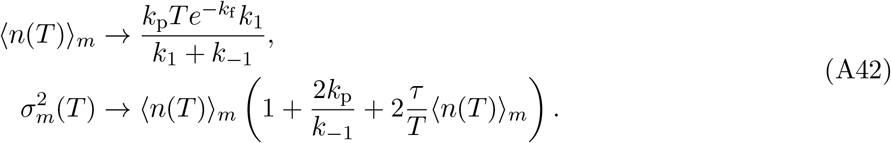

When *τ ≪ T* and *k*_p_ *→* 0, we have recovered the results in Eq. (A36) and Eq. (A40).

### A2 Information transmitted by KPR and Michaelis-Mentens schemes

In order to explain why the simplified KPR scheme performs better than the MM scheme, which is different only through a different choice of proofreading/processing time *τ*, we consider the DNA replication scenario.

We will provide a mathematical analysis as well as physical intuition from an information-theoretic perspective. To be specific, we consider the following two schemes:

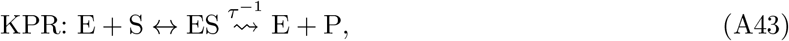

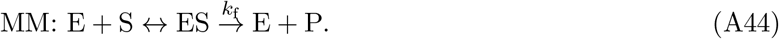

#### Mathematical analysis

In the scenario of DNA replication, we note that Scheme (A43) yields an error probability of

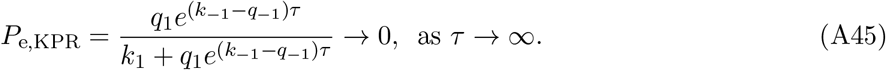

On the other hand, Scheme (A44) yields an error probability of

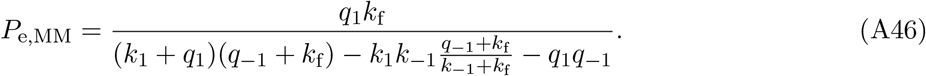

By investigating 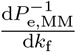, we find that *P*_e,MM_ is monotonically increasing with respect to *k*_f_, with the maximum 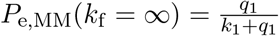 and minimum 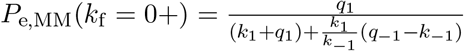.

To summarize, the error probability of Scheme (A43) converges to 0 at an exponentially fast rate, while the error probability of Scheme (A44) converges to a positive limit at a finite rate. In the no-proofreading limit, *k*_f_ = *∞* and *τ* = 0, both schemes yield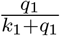, which depends only on the binding rate.

#### Physical intuition

Next, we provide an intuitive explanation of the above results from an information-theoretic perspective. To simplify the analysis, we consider the information yield of the two schemes *per binding-unbinding cycle*. We begin by introducing a timer *a* that records the time spent in the bound state (ES).

At time *t* = 0, we set the system in the bound state and set the timer *a* to 0. The unbinding waiting time *τ*_*−*1_ is exponential with rate *k*_*−*1_ or *q*_*−*1_, depending on the input *ξ* = 1 or *ξ* = 0. At the moment of unbinding, the timer *a* = *τ*_*−*1_ is read and used to determine whether the product is formed. In other words, this can be viewed as a two-step channel with the first step being the reading of the timer *a* and the second step being the determination of the product formation, which maps the timer value to the product formation.

The information yield of the first step is given by the mutual information between the input *ξ* and the timer (or age) *a*, in which we assume *ξ* is uniformly distributed. In the case of *k*_*−*1_ = 1 and *q*_*−*1_ = 2, the mutual information is given by ℐ(*ξ*; *a*) *≈* 0.078.

The second step involves a mapping from the timer value to the product formation. In the case of Scheme (A43), the mapping is deterministic with a threshold *τ*, given by 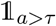. In the case of Scheme (A44), the mapping is stochastic with an additional exponential waiting time *τ*_f_ with rate *k*_f_, given by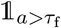. We can expect that because of the additional layer of stochasticity, the information yield of the MM scheme is much lower than that of the KPR scheme, as confirmed by the numerical results in Fig. A1. We also note that the information yield per binding-unbinding cycle is low. Thus, multiple rounds of binding and unbinding are required to achieve a high channel capacity.

**Fig A1.**
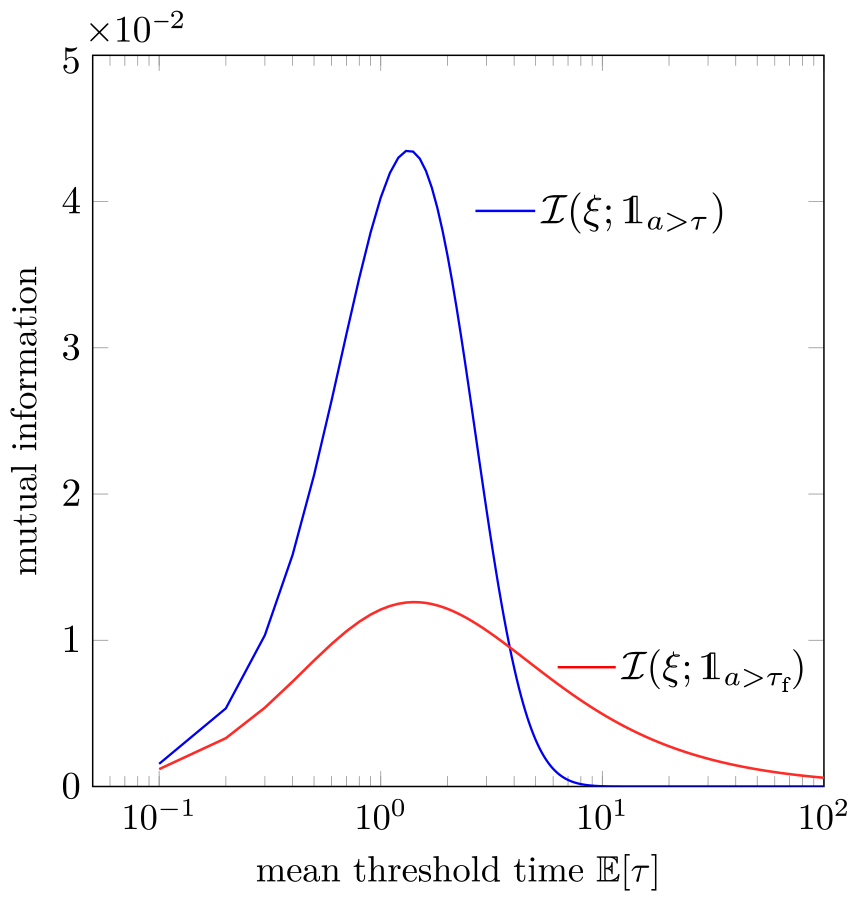
The mutual information between the input *ξ* and the final output for the KPR and MM schemes 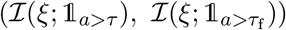 in a single unbinding event, for different mean proofreading time. We set *k*_*−*1_ = 1 and *q*_*−*1_ = 2.

### A3 Multistep binding model

In the main text, we have considered the analysis of the multistep limit of the classical KPR model, where the processing time from initial binding to full activation is taken to be a deterministic time *τ*. In the classical KPR model, the processing time is subject to noise due to finite number of steps in the activation process.

In this section, we fix the number of activation steps *m* = 6 and perform simulations using the same parameters as in the main text. The biophysical rationale derives from the six phosphorylation sites in the associated CD3 *ζ* chains. In particular, the phosphorylation kinetics of these sites are independent of the antigen types, except for two extra phosphorylation sites on ZAP70 that are regulated depending on the antigen types, which we do not explicitly consider in this paper. Figure A2 shows a qualitatively similar dependence of channel capacity on processing time *τ* as the deterministic limit in Fig. 6. The channel capacity of product concentration is maximized at *τ ≈* 0.5*/k*_1_, irrespective of *T*, while the optimal processing time for the channel capacity of first activation time increases with cell contact time *T*.

**Fig A2.**
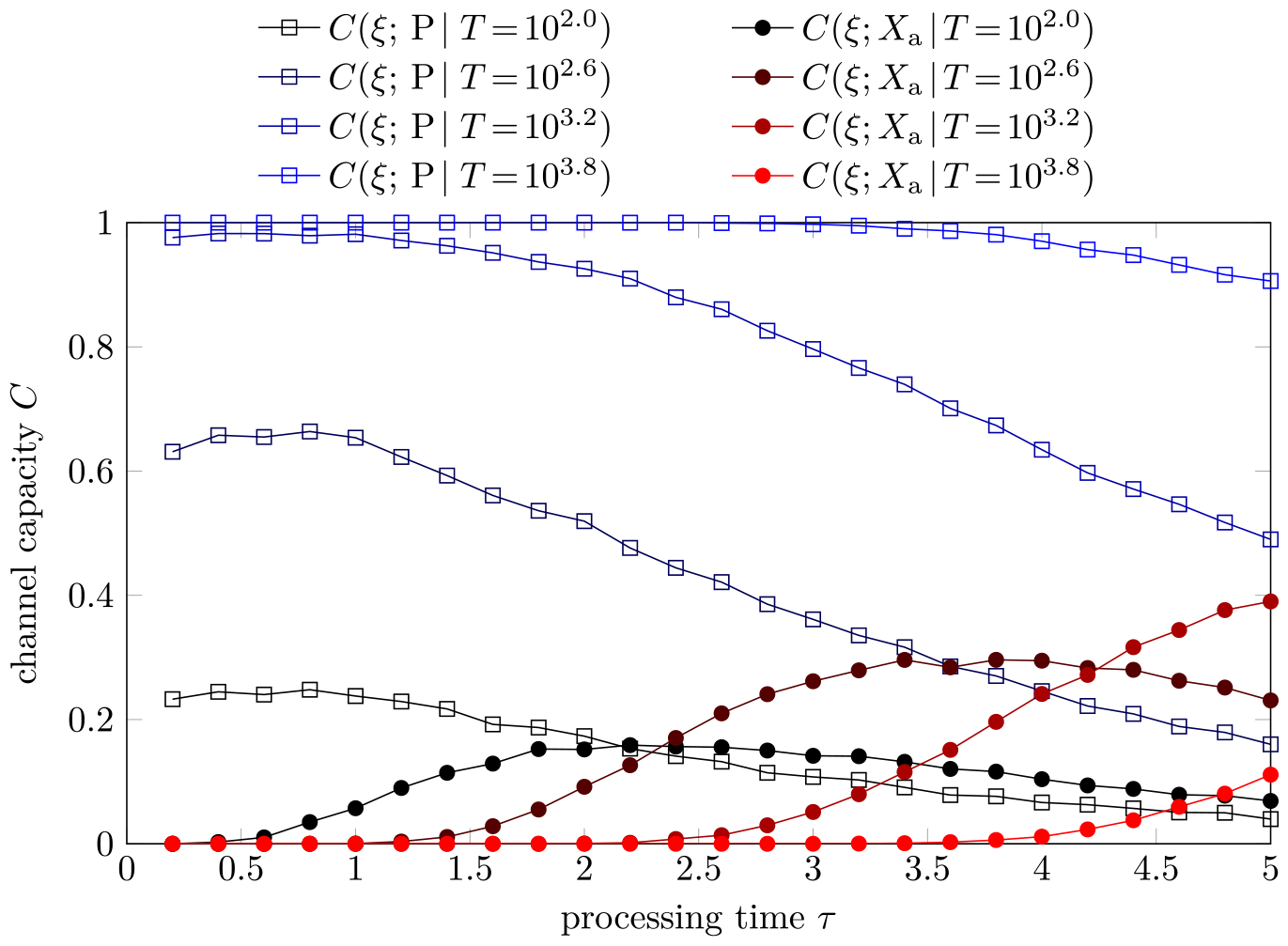
The channel capacities of product concentration (blue squares) and first activation times (red dots) as a function of processing time *τ* for various cell contact time *T* in a six-step multistep binding model. The parameter values are those used in Fig. 6.

Similar relationships are observed for the channel capacity of decision based on whether the protein number reaches a threshold P_th_ at time *T*, as shown in Fig. A3 compared to Fig. 9.

However, in both cases, the channel capacity of first-passage-time-based decision with 6-step activation is lower than that of the deterministic limit, which is a consequence of the noise in the activation process. The channel capacity of product concentration is similar to that of the deterministic limit, which may be explained by the buffering effect of product generation steps to upstream noises. In particular, decrease of *C*(*X*_th_ | P_th_) is dependent on P_th_. Larger P_th_ leads to a smaller decrease of channel capacity. The optimal processing time for first-passage-time-based discrimination also increases, compared to the deterministic limit. A quantitative analysis awaits further investigation.

**Fig A3.**
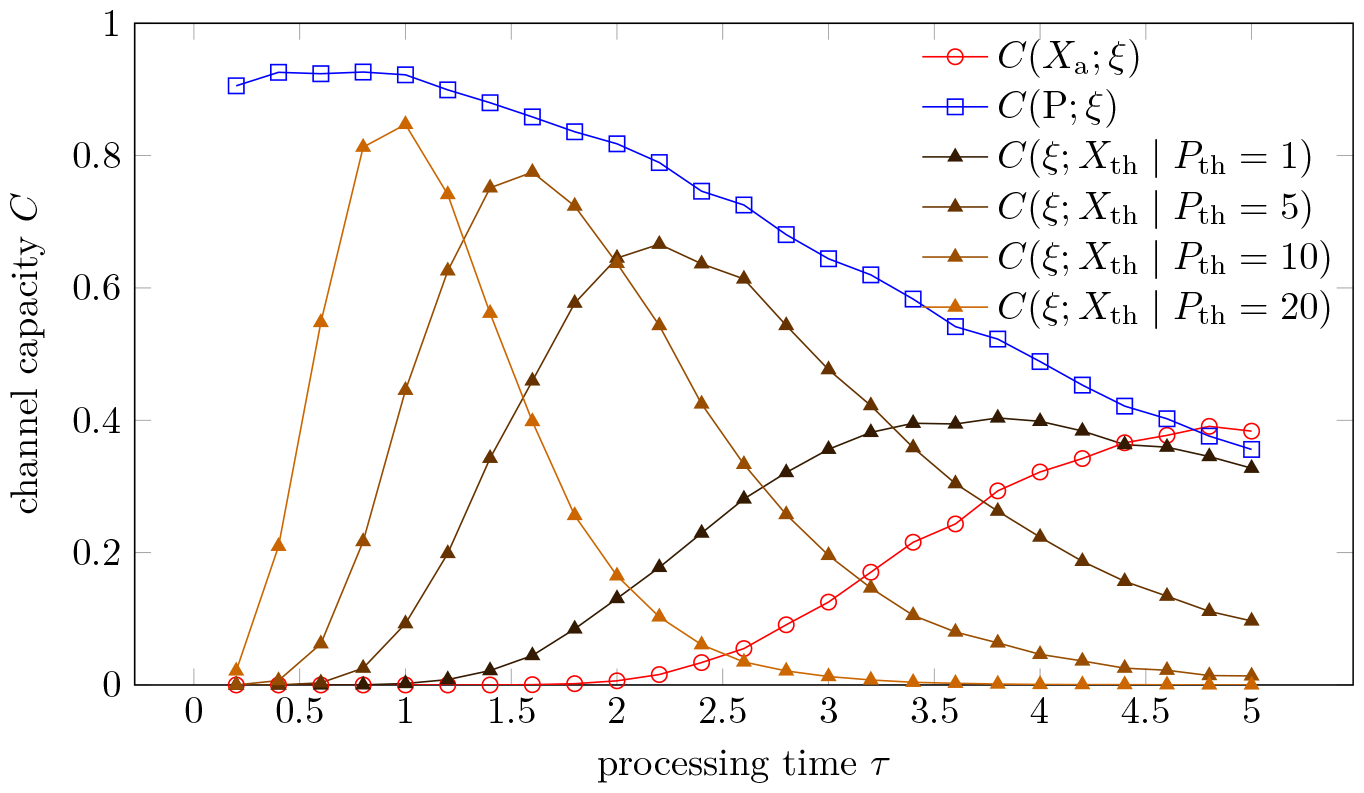
The channel capacity between the input *ξ* and the output *X*_a_, P(*T*), or *X*_th_ as a function of processing time *τ*. The parameters are kept to be the same as in Fig. 9. 10,000 independent Gillespie simulations are conducted for each *τ*.

### A4 Supplementary figures

**Fig S1.**
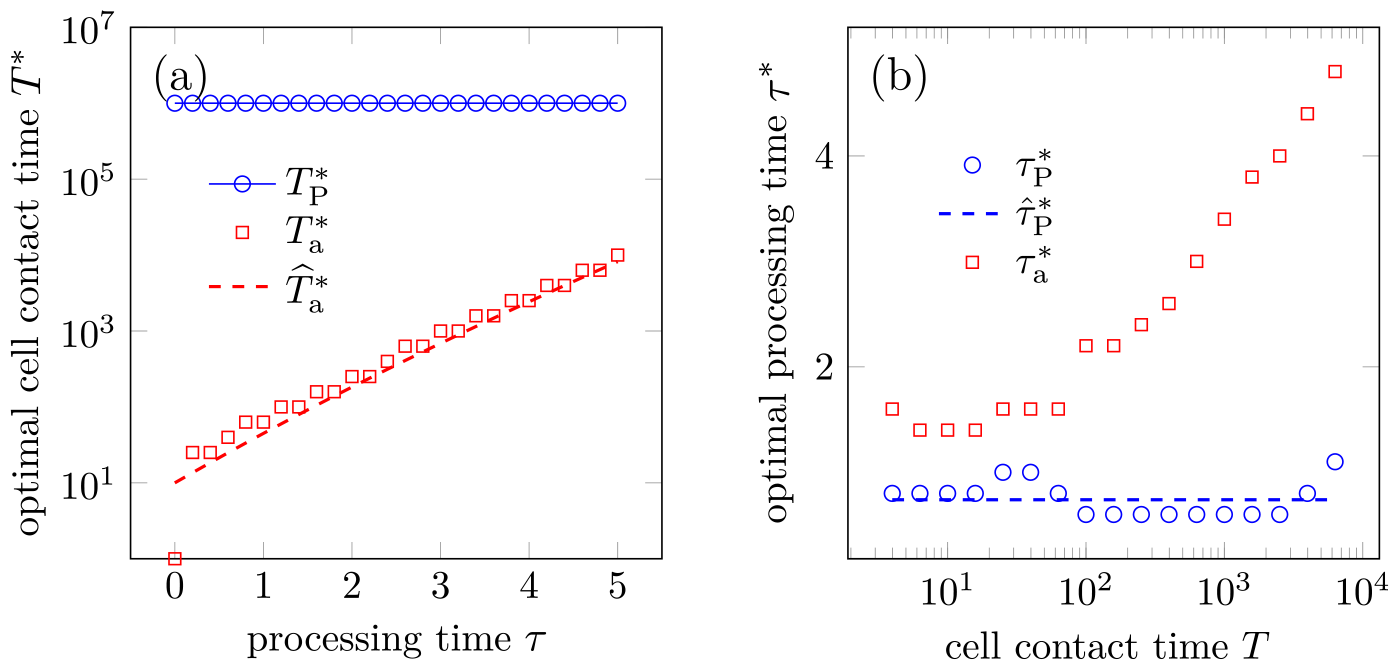
Dependence of optimal contact times and processing times on each other. Symbols represent results from simulations over a total time horizon of 10^6^, while the dashed curves represent analytic approximations. (a) Dependence of optimal cell contact times 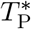 and 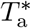 on processing time *τ* under product-based and FPT-based strategies. Since *C*(*ξ*; *P*) is nondecreasing with respect to *T*, the maximizing 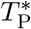 value is given as the upper limit of the simulation period. The approximation for 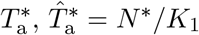, is given by Eq. (11). (b) Dependence of optimal processing times 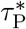 and 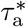 on cell contact time *T*. The analytic approximation 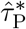 is given by Eq. (20). The parameters used are the same as those in Fig. 6.

**Fig S2.**
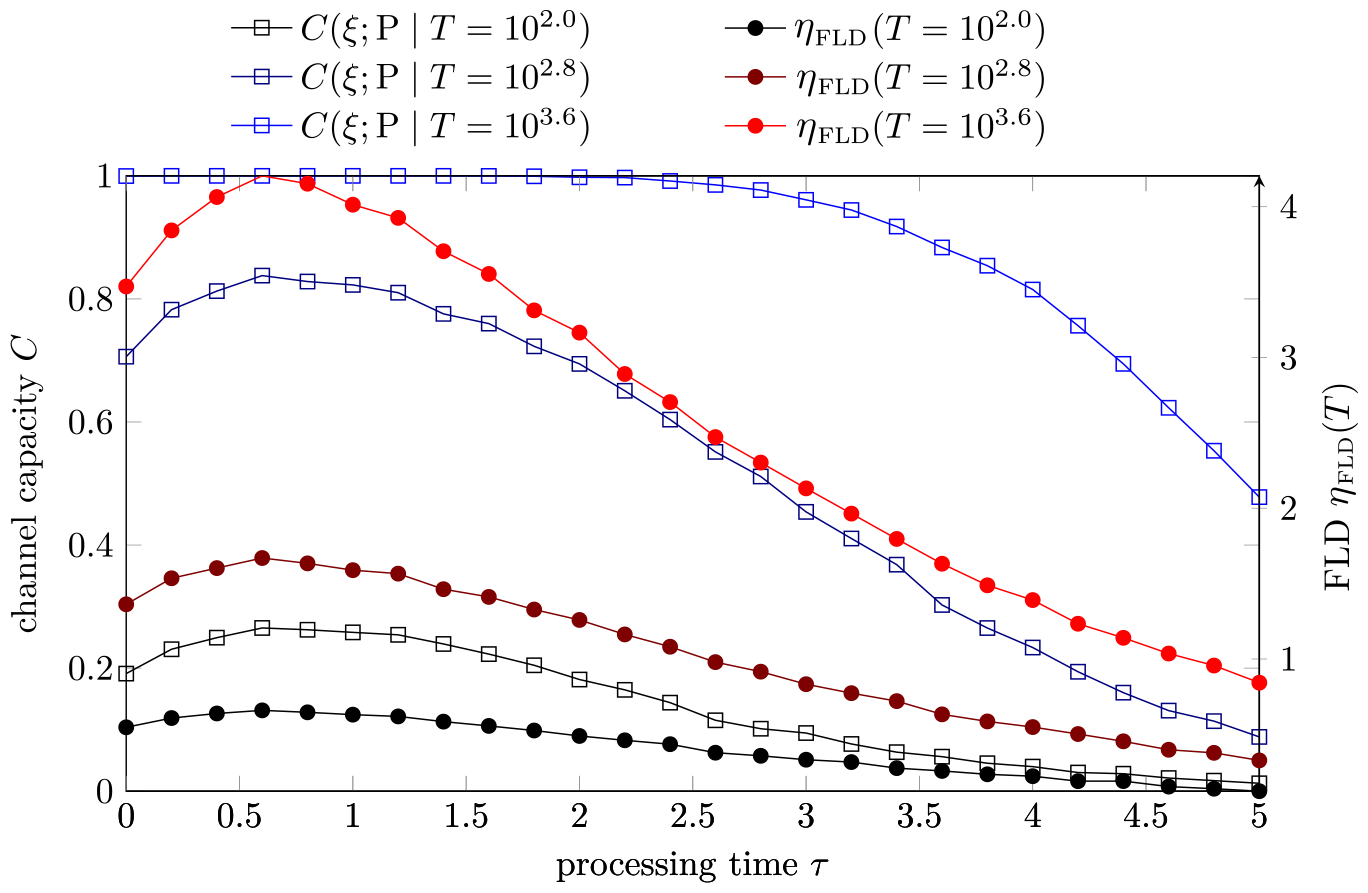
Dependence of the channel capacity between input *k*_*−*1_ or *q*_*−*1_ and the product P at time *T* and the Fisher linear discriminant on the processing time *τ* for different values of *T*. Both quantities are evaluated via numerical simulations from 10^4^ trajectories. We assumed 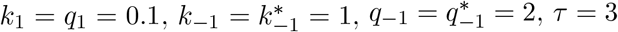, and *k*_p_ = 1 for a fast product formation rate.

**Fig S3.**
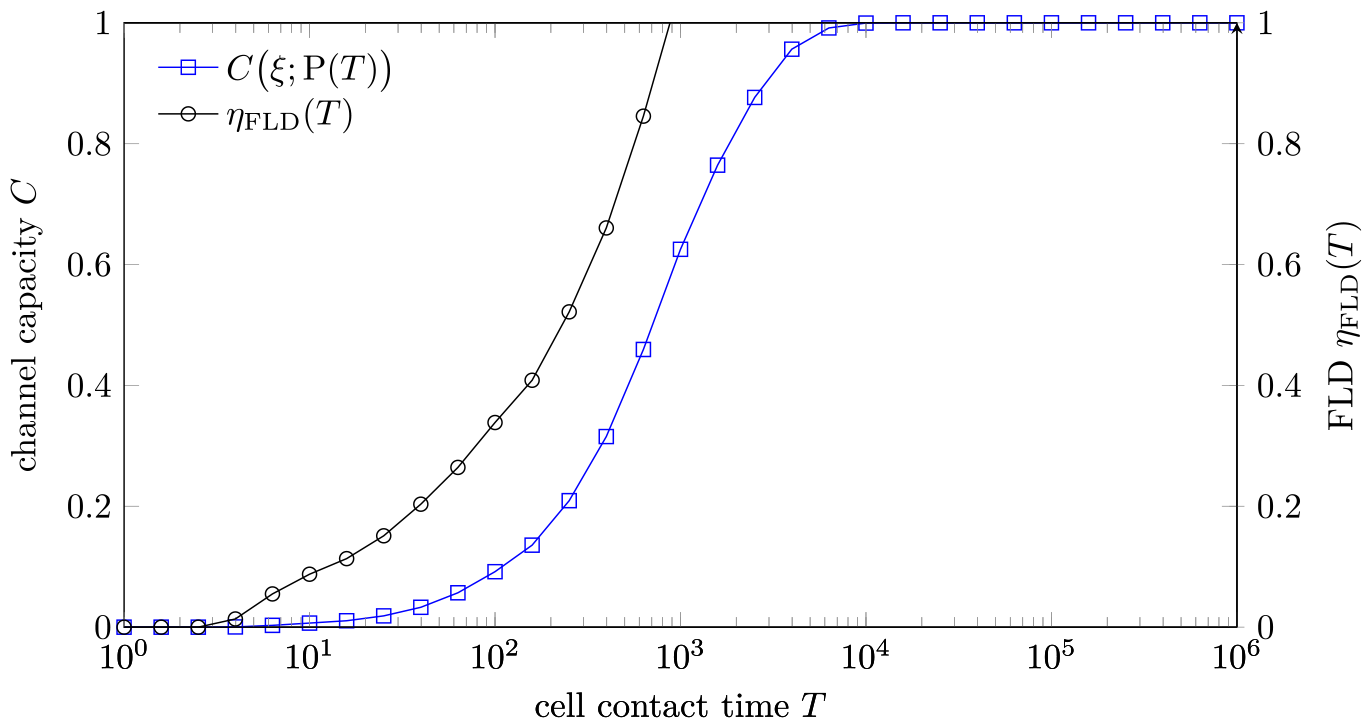
Dependence of the channel capacity between input *ξ* and the product P and the Fisher linear discriminant on the cell-cell contact time *T*. Both quantities are evaluated via numerical simulations from 10^4^ trajectories. We assumed 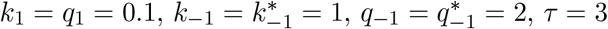, and *k*_p_ = 1 for a fast product formation rate. For *η*_FLD_, when *T >* 10^3^, their values are too high to be shown in the figure. We cropped the values of *η*_FLD_ to (0, 1) for better visualization.

**Fig S4.**
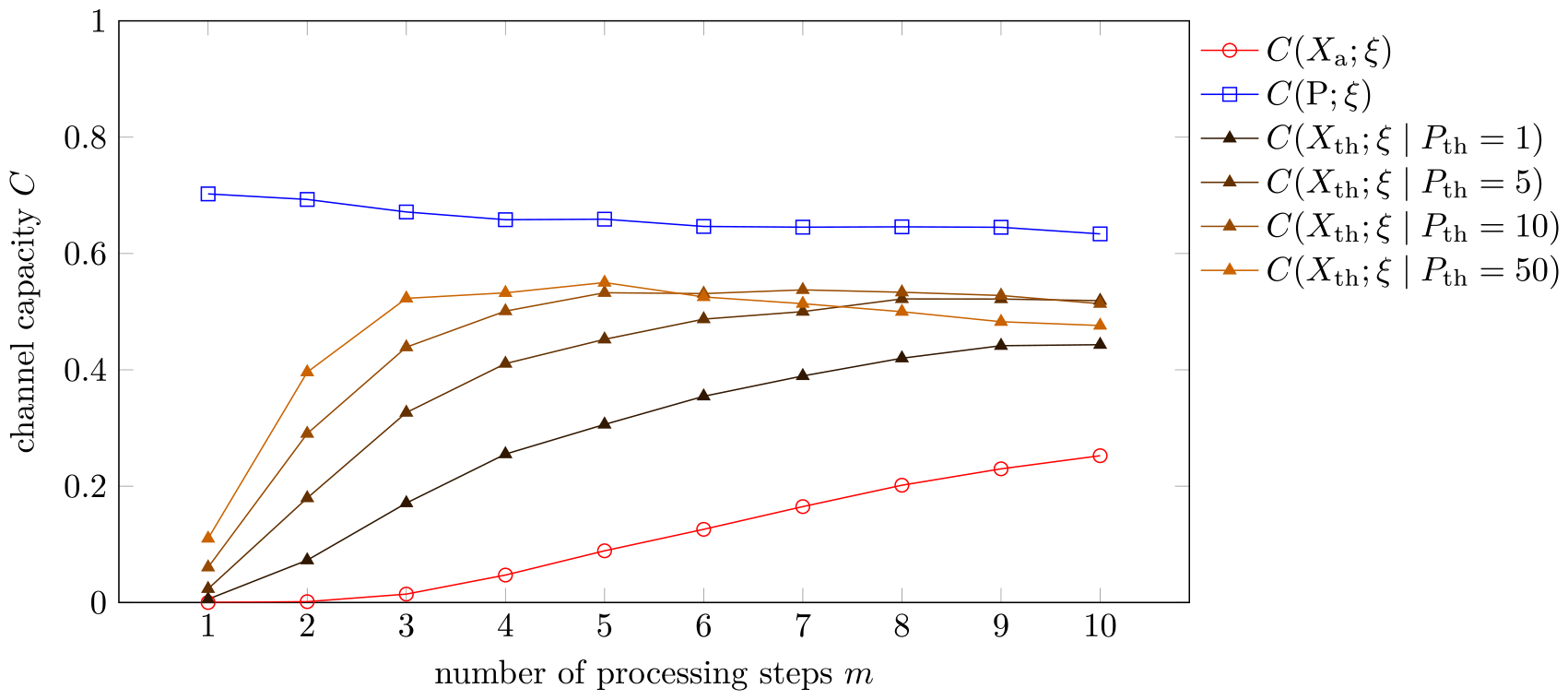
Dependence of the channel capacity between input *ξ* and different outputs on the number of proofreading steps *m*. They are obtained from numerical simulations of the multistep binding model with 10^5^ trajectories for each set of parameters. We assumed 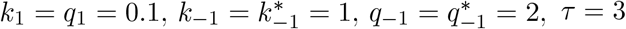, and *k*_p_ = 1. *T* is fixed at 1000. The cross-over between P_th_ = 50 and P_th_ = 10 is a result of finite sample stochasticity.

## Notes

### Competing Interest Statement

The authors have declared no competing interest.

